# Development of a High-Performance Open-Source 3D Bioprinter

**DOI:** 10.1101/2022.09.11.507416

**Authors:** Joshua W. Tashman, Daniel J. Shiwarski, Adam W. Feinberg

## Abstract

The application of 3D printing to biological research has provided the tissue engineering community with a method for organizing cells and biological materials into complex 3D structures. While many commercial bioprinting platforms exist, they are expensive, ranging from $5,000 to over $500,000. This high cost of entry prevents many labs from incorporating 3D bioprinting into their research. Due to the open-source nature of desktop plastic 3D printers, an alternative option has been to convert low-cost plastic printers into bioprinters. Several open-source modifications have been described, but there remains a need for a user-friendly, step-by-step guide for converting a thermoplastic printer into a bioprinter using components with validated performance. Here we convert a low-cost 3D printer, the FlashForge Finder, into a bioprinter using our Replistruder 4 syringe pump and the Duet3D Duet 2 WiFi for total cost of less than $900. We demonstrate that the accuracy of the bioprinter’s travel is better than 35 µm in all three axes, and quantify fidelity by printing square lattice collagen scaffolds with average errors less than 2%. We also show high fidelity reproduction of clinical-imaging data by printing a scaffold of a human ear using collagen bioink. Finally, to maximize accessibility and customizability, all components we have designed for the bioprinter conversion are provided as open-source 3D models, along with instructions for further modifying the bioprinter for additional use cases, resulting in a comprehensive guide for the bioprinting field.

## Introduction

Additive manufacturing has disrupted multiple industries because it enables the fabrication of complex 3D parts, rapid design iteration, low-cost customization, and the use of a growing range of engineering-grade materials (*1*). This transition has been supported by researchers developing new 3D printing methodologies and companies producing industrial-scale 3D printers for selective-last sintering (SLS), digital light processing (DLP), stereolithography (SLA), binder jetting, and thermoplastic filament extrusion. 3D bioprinting has the potential to bring similar improvements to the tissue engineering field by building cellularized constructs, and ultimately functional tissues and organs (*2*–*5*). Instead of polymers, metals, or ceramics, in 3D bioprinting it is the bioink that is printed, where the term bioink as used here includes high-density cell slurries, synthetic and natural hydrogels, cell-laden hydrogels, biomaterial inks, and combinations thereof. However, because 3D bioprinting is still primarily at the research and development stage, any barriers to widespread adoption serve to limit innovation. Chief among these barriers is the high cost of commercial, research-grade 3D bioprinting platforms, which range from $10,000 to over $1,000,000. At these price points, a capital equipment purchase and dedicated funds are typically required, which limits access to core facilities and well-funded research laboratories. Furthermore, the hardware and software of many of these 3D bioprinting platforms are difficult to modify for custom applications without incurring additional cost, have limited compatibility with new biomaterials, and use proprietary printing software and a closed hardware ecosystem.

A solution for these issues emerged with the open-source 3D printing community that began in the early 2000s and accelerated with the expiration of national and international patents on fused deposition modeling (FDM) in 2009 (*6*). For the first time, plastic 3D printing went from a relatively expensive technique, using proprietary equipment and materials dominated by large companies, to an open-source movement spurred by startup companies and inexpensive 3D printers that could be used by anyone. As early as 2012, researchers began to convert these low-cost thermoplastic printers, which were continuously being improved by the open-source community, into bioprinters capable of producing high-quality results for tens of thousands of dollars less than commercial alternatives. Similarly, early work on custom-built bioprinters like the fab@home project at Cornell showed the potential of building open-source platforms at relatively low cost (*7*). Over this period our research group converted a wide range of open-source thermoplastic printers (e.g., MakerBot Replicator, LulzBot Mini 2, PrintrBot Simple Metal, FlashForge Creator Pro, MakerGear M2) into high-performance 3D bioprinters (*8*–*10*). This has enabled us to leverage the high-quality 3-axis motion system that these open-source printers already have while only needing to add the components, such as the syringe-pump extruder, specifically required for bioprinting cells and liquid bioinks. Further, our approach uses the same stepper motor from the original filament extruder of the thermoplastic printer to drive the syringe-pump extruder of the bioprinter. This means multiple high quality open-source software packages can be used to slice 3D models into G-code and to control the printing process, just as in plastic printing.

While there have been several publications describing 3D bioprinter modifications, including extruders we have designed (*9, 11*), as a whole the field lacks a comprehensive guide for building a complete, customizable open-source 3D bioprinter platform using validated and tested components (*12*–*17*). Here we describe the modification of a, low-cost thermoplastic 3D printer that is widely available, into a sub $1000 bioprinter. Since 2018, we have also run an internationally attended open-source 3D bioprinting workshop at Carnegie Mellon University where participants build their own bioprinter, learn how to use them for FRESH 3D bioprinting, and then take the bioprinters back to their home institutions for future research. These efforts have served to validate our bioprinter designs and modifications and step-by-step guides for a range user backgrounds and experience levels, and have produced multiple high-impact publications (*18*–*24*). While we use a FlashForge Finder as the printer here, the approach we describe is readily adaptable to nearly any low-cost and open-source printer on the market. To do this we have created instructions requiring minimal knowledge of electronics or mechanical fabrication, and use previously published open-source printer components and syringe pump extruders capable of producing high quality printed constructs (*11*). To ensure easy adoption and future customizability, we replace the proprietary motion control circuit board of the Finder with the Duet 2 WiFi, an open-source, highly adaptable, easy to use, and very well documented motion control board from Duet3D. We also outfit the printer with our Replistruder 4 open-source syringe pump, which builds on almost a decade of designs from our lab, and which has been used in multiple published studies (*9, 11, 18*). The end result is a low-cost 3D bioprinter with performance on par or better than commercial alternatives and with a high degree of hardware and software customizability that is critical to printing new bioinks and developing advanced applications.

## Results

### Converting a Plastic Printer into a Bioprinter

When converting a plastic 3D printer into a bioprinter there are a sequence of steps that generally occur in the same order (Fig. 1). First, the electronics and control system of the plastic printer either need to be adapted to bioprinting through modification, or they need to be replaced with an alternative. The proprietary FlashForge Finder motion control circuit board (Fig. 2A, green rectangle) is replaced with the open-source Duet 2 WiFi motion control circuit board (Fig. 2B, blue rectangle). This is done to improve the motion control performance, provide WiFi access, and to facilitate rapid firmware customization through the Duet web-based interface without needing to use additional software. The step-by-step instructions for this process are laid out for the FlashForge Finder, and can be adapted for most desktop 3D printers (details provided in the supplemental assembly guide, and Supplementary Figs. S1-S15). Next, the thermoplastic extruder that came with the printer is replaced with the Replistruder 4, an open-source, high-performance syringe pump extruder we have previously developed (*11*). Most parts of the Replistruder 4 are readily 3D printed out of plastic and assembled using commonly available hardware (Fig. 2C). A carriage platform was designed to fit on the existing linear motion components of the printer and provide a mounting point for the Replistruder 4. This X-axis carriage has pockets for the bearings already mounted on the X-axis linear rails as well as channels for routing and retaining the X-axis belt that drives motion along the axis. Additionally, four mounting points with recessed M3 hex nuts are incorporated to allow the Replistruder 4 to be attached to the X-axis carriage (Fig. 2D). The thermoplastic printhead that comes pre-installed on the Finder is replaced with the X-axis carriage/Replistruder 4 assembly (Fig. 2E, F, details in supplemental assembly guide, and Supplementary Figs. S16-S23). At the completion of these steps, the Replistruder 4 (Fig. 2G, blue arrow) is mounted on the X-axis of the printer in the X-axis carriage (Fig. 2G yellow arrow), and the motors are connected to the Duet 2 WiFi, which is positioned in the back cabinet of the bioprinter (Fig. 2H, green arrow). With these modifications the FlashForge Finder is transformed into an open-source bioprinter with a high-performance extruder and motion control system.

**Figure 1:**
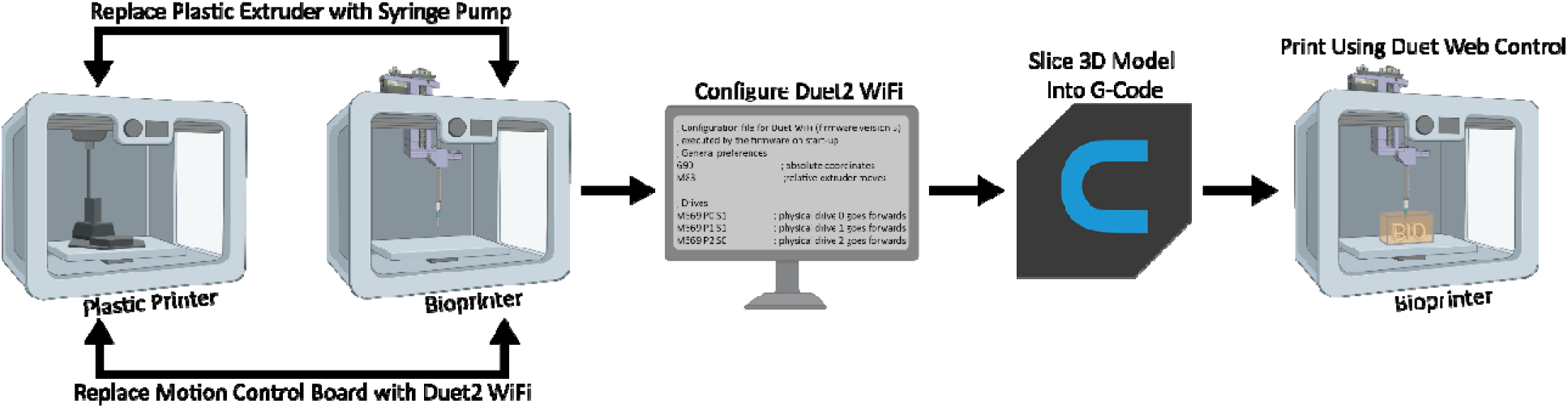
Steps in Converting a Plastic Printer to A Bioprinter. The plastic printhead and the control system of the plastic printer are switched to a syringe pump and Duet2 WiFi. The Duet2 WiFi is then configured to run a bioprinter. To bioprint, the desired 3D model is sliced into machine pathing (G-code) and then Duet Web Control executes the print using the desired bioink.

**Figure 2:**
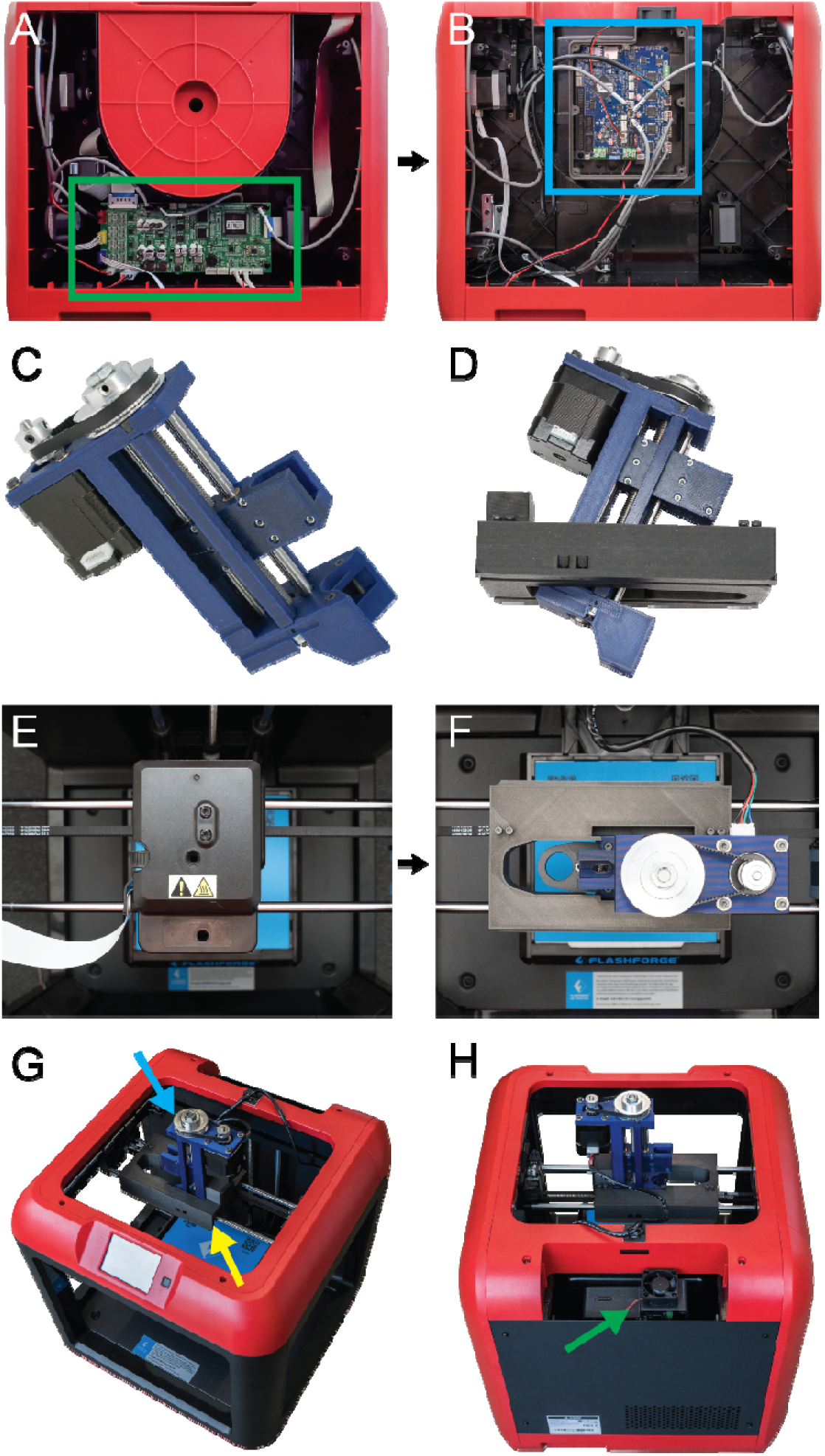
Converting the FlashForge Finder into a Bioprinter. **(A)** The original control electronics and wiring (green rectangle). **(B)** Controls replaced and adapted to the Duet 2 WiFi (blue rectangle). **(C)** The Replistruder 4 is printed and assembled. **(D)** The X-axis carriage for the FlashForge Finder is printed and the Replistruder 4 is mounted to it. **(E)** Top-down view of the Finder’s plastic printhead that is to be removed. **(F)** Top-down view of the Replistruder 4 after it is mounted in the printer. **(G)** Additional view showing that the plastic printhead has been replaced with a Replistruder 4 syringe pump (blue arrow) mounted to a custom X-axis carriage (yellow arrow). **(H)** The Duet 2 WiFi is mounted in a 3D printed case covered in the back cabinet of the Finder (green arrow).

The Duet 2 WiFi provides several key advantages over the stock motion control circuit boards found in the FlashForge Finder and other low-cost desktop 3D printers. First, the Duet’s WiFi based web interface allows for easy, in-browser access to printer movement, file storage and transfer, configuration settings, and firmware updates. This is in contrast to most 3D printers, where the process of editing configuration settings requires flashing the motion control board firmware using 3^rd^ party software. This can be challenging and intimidating for an inexperienced user and may lead to accidental changes or firmware corruption. Second, the Duet 2 adds many advanced motion control improvements including (i) a 32-bit electronic controller, (ii) high performance Trinamic TMC2260 stepper controllers, (iii) improved motion control with up to 256x microstepping for 5 axes, (iv) high motor current output of 2.8A to generate higher power, (v) an onboard microSD card reader for firmware storage and file transfer, and (vi) expansion boards adding compatibility for 5 additional axes, servo controllers, extruder heaters, up to 16 extra I/O connections, and support for a Raspberry Pi single board computer. Together, these features provide high precision motion control and extensive expandability with an easy-to-use web interface that enables rapid customization and improved performance beyond standard desktop plastic printers.

### 3D Bioprinter Mechanical Performance

After conversion, the X, Y, and Z axis travel limits were measured in order to determine the build volume of the 3D bioprinter. For the X-axis, travel was 105 mm, for the Y-axis travel was 150 mm, and for the Z-axis travel was 50 mm, resulting in an overall build volume of 787.5 cm^3^ (Fig. 3A). For a stepper motor driven motion system, such as this 3D bioprinter and most commercial 3D printers, the most important control parameter affecting performance is the steps per mm calibration for each of the three axes. This number determines how many pulses, or steps, must be sent to the stepper motors that drive each axis to move them precisely one millimeter each. For the X and Y axes, which are belt driven, the formula for this is *steps/mm = (steps/rotation * microsteps)/(belt pitch * pully teeth)*. For the Finder these parameters are the nominal pitch of the driving belt (2 mm), the number of teeth in the motor’s pully (17 teeth), the number of steps in a full rotation for the motor (200 steps), and the number of microsteps that the Duet 2 WiFi interpolates between the full steps (set to 64 microsteps). In this case the nominal steps/mm for the X and Y-axis is 376.5. For the Z-axis, which uses a leadscrew, the formula is *steps/mm = (steps/rotation * microsteps)/(screw pitch * screw starts)*. The finder uses a 4 start, 2 mm pitch lead screw so the nominal steps/mm for 16x microstepping is 400 steps/mm.

**Figure 3:**
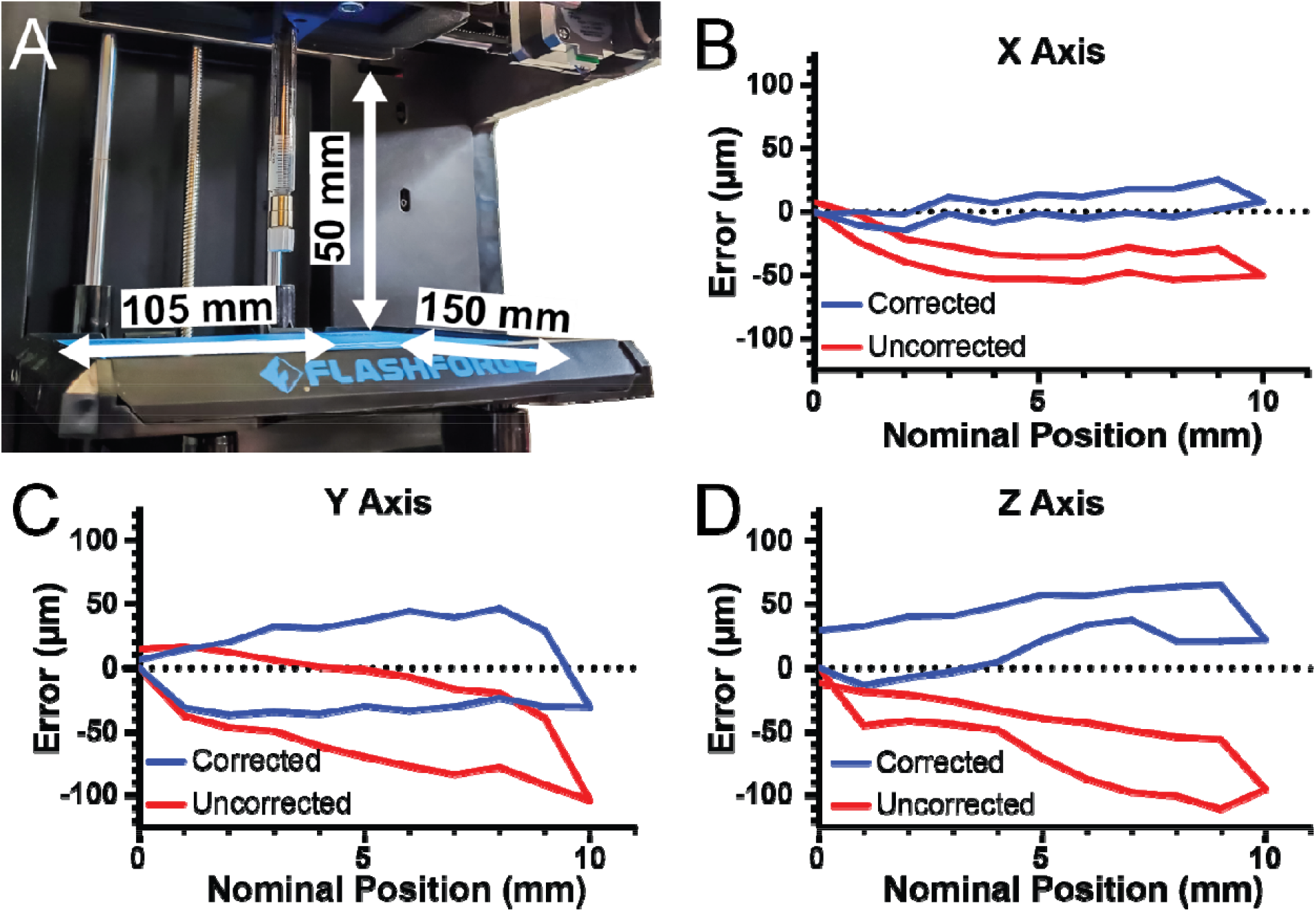
Measurement and Correction of Printer Travel. **(A)** The X-axis travel is 105 mm, the Y-axis travel is 150 mm, and the Z-axis travel is 50 mm. **(B)** The error of travel for the X-axis across a 10 mm window before correction (red) and after correction (blue). **(C)** The error of travel for the Y-axis across a 10 mm window before correction (red) and after correction (blue). **(D)** The error of travel for the Z-axis across a 10 mm window before correction (red) and after correction (blue).

When comparing 3D bioprinter performance there are several key specifications to consider that relate directly to print quality. Most thermoplastic 3D printers provide specifications for resolution, which is defined as the smallest step the printer can take in any direction. The reported numbers for the FlashForge Finder are 11 µm in XY and 2.5 µm in Z. This is on par with many commercial bioprinting platforms such as the CellInk BioX (1 µm in XYZ), LulzBot Bio (10 µm in XY and 5 µm in Z), and Allevi 2 (7.5 µm in XY and 5 µm in Z). Additional specifications related to the motion system are positional error, which is defined as the absolute deviation of the current location of the printhead from the intended location, and repeatability, which is defined as the maximum absolute deviation in measured position from the average measured position when attempting to reach that position multiple times. These more sophisticated metrics are largely absent in the bioprinting space and can vary between individual printers based on mechanical components and accuracy of assembly. Additionally, the resolution provided in bioprinter specifications are generally ideals based off of the nominal dimensions of the gears, pulleys, and screws used to assemble the motion system. None of the previously mentioned manufactures provide measurements of actual resolution, that is error across the full distance of travel, or repeatability. These measurements are commonly provided with ultra-high-end motion platforms such as those from Aerotech and Physik Instrumente (*25, 26*). To determine and then optimize the real world performance of these low cost 3D printer based systems it is necessary to measure the travel with an external tool.

To verify that the nominal steps/mm values were correct, we quantified positional error of our system along each axis near the center of travel with 2 µm precision. For the X-axis, there was a systematic under-travel using the nominal steps/mm (Fig. 3B, red curve). Using the maximum error at 10 mm of travel we determined the number of missed steps per mm and corrected the value, and with this corrected steps/mm the average travel error over the 10 mm window was 7.9 µm (Fig. 3B, blue curve). For the Y-axis there was systematic under-travel using the nominal steps/mm (Fig. 3C, red curve) and after correction this was reduced to 29.1 µm (Fig. 3C, blue curve). Finally, for the Z-axis there was systematic under-travel using the nominal steps/mm (Fig. 3D, red curve) and after correction was reduced to 32.3 µm (Fig. 3D, blue curve). The values also enable calculation of the unidirectional repeatability, which is the accuracy of returning to a specific position from only one side of the axis (e.g., from 0 to 5 mm) and the bidirectional repeatability, which is the accuracy of returning to a specific position from both sides of the axis (e.g., from 0 to 5 mm and from 10 to 5 mm). For the X-axis, the unidirectional repeatability was 3.9 µm, and the bidirectional repeatability was 16.4 µm. For the Y-axis, the unidirectional repeatability was 11.5 µm, and the bidirectional repeatability was 63.9 µm. For the Z-axis, the unidirectional repeatability was 8.7 µm, and the bidirectional repeatability was 38.7 µm. Together these measurements demonstrate that with calibration the travel of our converted bioprinter had a maximal error of 35 µm and repeatability in worst case situations of 65 µm. Whereas before calibration there was a linearly increasing error in position, afterwards this error is significantly decreased. Without this calibration, or at least measurement of the errors, it would be impossible to determine if flaws in printed constructs were due to the printer itself or other factors impacting print quality.

### Assessing Bioprinter Printing Fidelity

Feature resolution of printed bioinks is typically not quantified for 3D bioprinters because they cannot print biological material in a manner approaching the mechanical limitations of the systems. However, with the recent advancements made in embedded bioprinting techniques such as FRESH (*10*), it is now possible to perform extrusion bioprinting with resolutions approaching 20 µm. To demonstrate bioprinter printing performance we generated a square-lattice scaffold design consisting of 1,000 and 500 µm filament spacing (Fig. 4A) to measure accuracy when FRESH printed from a collagen type I bioink (Fig. 4B). To measure the grid spacing we captured a 3D volumetric image using optical coherence tomography (OCT) (Fig. 4C) (*27*), which revealed close agreement between the as-designed and measured dimensions (Fig. 4D). This was followed by a more complex design based on a 3D scan of an adult human ear (Fig. 4E). This model was printed using collagen (Fig. 4F). To analyze the accuracy we captured a 3D volumetric image of the printed ear using OCT (Fig. 4G) (*27*). The 3D reconstruction demonstrates recapitulation of the features of the model and the gauging quantification revealed a deviation of -29 ± 107 µm (mean ± STD) between the FRESH printed ear and the original 3D model (Fig. 4H). Together, these two examples demonstrate that the average error and standard deviation of the printed scaffolds are within the mechanical limitations of the bioprinter we built, and is on par with commercial bioprinters (*28*).

**Figure 4:**
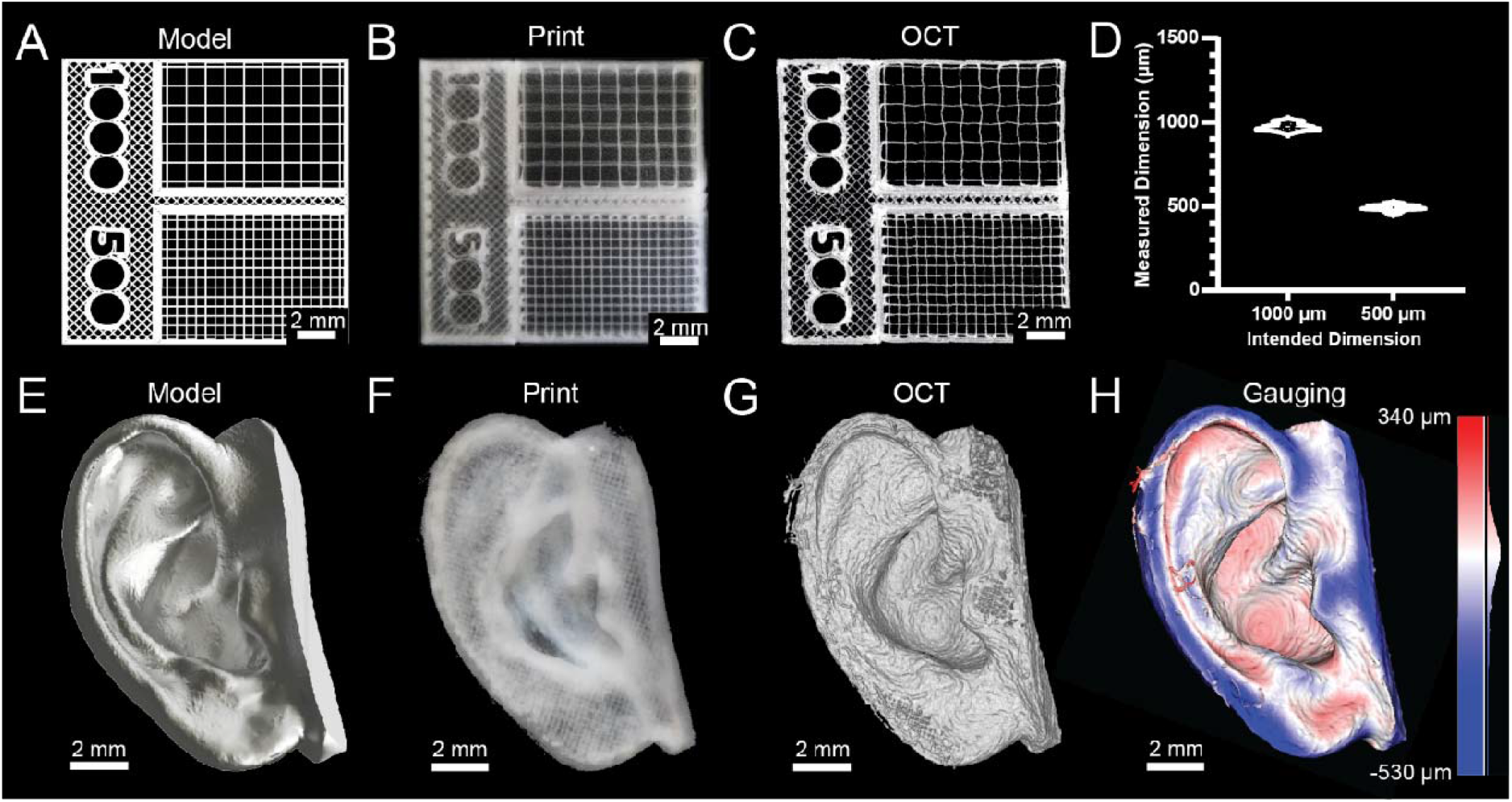
Printing Dimensionally Accurate Grids and Organic Shapes. **(A)** Model of a gridded scaffold with 500 µm and 1000 µm grids. **(B)** Photograph of the gridded scaffold printed in collagen type I. **(C)** An OCT image of the gridded scaffold print. **(D)** Analysis of the accuracy of the gridded scaffold print (mean ± STD.; n = 11 measurements for 1000 µm grid, n = 26 measurements for 500 µm grid, p<.0001 [****] by Student’s two-tailed, unpaired t-test). **(E)** A 3D model of an ear. **(F)** A photograph of the ear printed in collagen type I. **(G)** An OCT volumetric image of the printed ear. **(H)** Quantitative gauging of the ear print against the original 3D model.

## Discussion

Here we have designed and built an open-source 3D bioprinter and provided detailed instructions using the FlashForge Finder as an example for the 3D printer conversion. While several bioprinter conversions have been published and have provided useful insight (*12*–*17*), there has remained a need for a user-friendly, step-by-step conversion guide using components that are widely available and that have been validated to perform with high level of performance. The FlashForge Finder is low cost (∼$300) and widely available around the world from online retailers, ensuring it will be accessible broadly to the research community. Additionally, we use a well-supported, open-source 3D printer motion control circuit board, the Duet 2 WiFi from Duet3D. All the components required for the conversion are either 3D printed in plastic such as PLA or widely available basic fastening, linear motion, and motor hardware. The total cost, including the Replistruder 4 components, is less than $900. Furthermore, while the step-by-step guide (see supplemental material) provides specific instructions for the FlashForge Finder, the instructions are generalizable and can be used to convert a wide array of other 3D printers into bioprinters. Together, these elements make our conversion both an easy-to-use introduction for a novice and a starting point for more advanced printer conversions and customizations.

Essential to the 3D bioprinting process is ensuring that the object being printed matches the dimensions and geometry of the input 3D CAD model. For the Replistruder 4 syringe pump extruder, this verification has been performed in previously published work (*11*). For the 3D bioprinter built here, nominal values for motion control parameters produced a maximum error over a 10 mm travel range of ∼100 µm, or ∼1% (Figs. 3B-D). For many bioprinting applications this is adequate accuracy, however, we show that by further calibrating the printer the accuracy can be improved three-fold to ∼33 µm (Figs. 3B-D). Considering that most bioprinter nozzle diameters range from 100 to 500 µm, this will produce high quality results for a wide range of applications (*9, 10, 27*). While analysis of the motion itself is important, validation of fidelity is best established by 16easurement of constructs 3D bioprinted with the platform. For the square lattice scaffold the 1,000 and 500 µm grids matched their intended dimensions closely and the slight deviations are thought to be due to post-print handling of the 90 µm diameter, 1000 µm long collagen filaments. In line with this reasoning, the 500 µm grid, which differed only by having more frequent contact points between orthogonal filaments, had improved accuracy compared to the 1000 µm grid. The ear scaffold showed that the bioprinter can produce more complex 3D shapes and the quantitative gauging analysis confirmed that the fidelity achieved using collagen bioink was similar to our published results using other FRESH 3D bioprinters (*10, 11, 29*). These examples demonstrate that the converted 3D bioprinter is capable of both accurate motion and high-fidelity printing of the intended geometry.

An important aspect of open-source hardware, aside from its lower cost, is the ability to modify and expand its capabilities for specific applications. For the previously published Replistruder 4 extruder, we provided complete print and design files that are easily modified in standard CAD software. Here we build upon this by providing complete print and design files for the X-axis carriage and various sample holders, including for 35 mm Petri dishes and multi-well plates. These files enable modification of the printer for various syringe types and sizes, as well as for various print containers. The Duet 2 WiFi allows for expanded capabilities as well, including heaters, chillers, and sensors that can be controlled using its advanced implementation of G-code. Additional tools are also supported such as multiple syringe pump extruders, lasers, subtractive tools (e.g., mills), UV light switching, automated tool switching, and G-code macros. Together this broad potential for extensibility and modification makes our 3D bioprinter a powerful platform for the development of advanced 3D bioprinting applications.

Finally, the 3D bioprinter we have developed here is the result of multiple years of development, during which we worked with engineers, scientists and physicians, and trained them to build and use these systems. It is based on the “3D Bioprinting Open-Source Workshop” that we have organized at Carnegie Mellon University since 2018, and where we have taught trainees from research labs across the world, including Australia, Canada, Israel, Japan, Korea, and the United States. The conversion of the FlashForge Finder into a 3D bioprinter described here was used for the 2021 workshop and served as the basis for the included step-by-step conversion guide (see supporting material). Through this publication we hope to expand the number researchers that can learn and benefit from this effort. Further, we hope that this 3D bioprinter can serve as an example of the impact that open-science can have on accelerating research advances and the importance of putting low-cost and high-performance scientific tools in the hands and labs of as many researchers as possible.

## Materials and Methods

### Measurement and Calibration of Travel

To measure travel of the printer with high precision we utilized a Mitutoyo Absolute Digimatic Indicator (Mitutoyo, Japan, 542-500B) with ±2 µm accuracy mounted to a Noga magnetic base for alignment with the axes of the printer. To measure a given axis the gauge was aligned parallel to the axis and then run to full travel then backed off to 100 µm less than its full travel using the printer controls on the Duet Web Control (to ensure any backlash on the initial travel was taken up). The axis was then moved away from the gauge in 1 mm nominal increments using the Duet Web Control. The actual position, taken from the gauge, was recorded. This process was repeated until a 10 mm nominal travel was complete, then the process was reversed until the nominal position was 0 again. The zero position was reset, and the process was repeated two more times. From these measurements the error was defined as the absolute difference between the nominal position and the measured position. The unidirectional repeatability was the error for the second and third trials when returning to each of the positions in each direction when compared to the first trial. The bidirectional repeatability was the absolute value of the difference between the actual measurements of position at the paired nominal locations (moving away and then returning to zero).

To calibrate the axes the nominal steps/mm for each axis is first input into the Duet WiFi 2 configuration file. The maximal error at 10 mm travel from the three trials was determined, and then the steps/mm for each axis was scaled proportionately to the error. That is if the nominal steps/mm was 376.5 and there was a total travel of 9.887 mm the correction would be (10 mm)/(9.887 mm)*(376.5 steps/mm) = 380.8 steps/mm. This process was then repeated for the other two axes. After calibration, the measurement process can be repeated to verify the accuracy.

### 3D Models of Printed Parts and Scaffolds for Bioprinter Conversion

All 3D CAD models for printed plastic components of the FlashForge Finder conversion and the Replistruder 4 were generated using Autodesk Inventor 2020 (Autodesk). The CAD files and STL files for the Replistruder 4 can be downloaded from Zenodo at https://doi.org/10.5281/zenodo.4119127. CAD and STL files can be downloaded from Zenodo at https://doi.org/10.5281/zenodo.7067499 for the FlashForge Finder. The grid model printed with 500 and 1000 µm grid spacing was generated using Autodesk Inventor 2020. The ear model was purchased from http://www.cgtrader.com/ and was generated by seller Sakura-pms.

### Generation of G-code

The G-code for printed plastic components of the Flashforge Finder conversion and the Replistruder 4 were generated using PrusaSlicer (Prusa). All models were printed from PLA plastic at 60% infill successfully without support material. The G-code for the ear model was generated using Cura 4.3.0 (Ultimaker). The G-code for the grid model was generated using Simplify 3D (Simplify3D).

### FRESH Gelatin Microparticle Support Bath and Generation

FRESH v2.0 gelatin microparticle support bath was generated via a complex coacervation method to produce gelatin microparticles, based on published methods (*10*). Briefly, 2.0% (w/v) gelatin Type B (Fisher Chemical), 0.25% (w/v) Pluronic® F-127 (Sigma-Aldrich) and 0.1% (w/v) gum Arabic (Sigma-Aldrich) were dissolved in a 50% (v/v) ethanol solution at 45ºC in a 1 L beaker and adjusted to 7.5 pH by addition of 1M hydrochloric acid. An overhead stirrer (IKA, Model RW20) was then used to maintain mixing while the beaker was sealed with parafilm to minimize evaporation, and the mixture cooled to room temperature with stirring overnight. The resulting solution was transferred into 250 mL containers and centrifuged at 300 g for 2 minutes to compact the gelatin microparticles. The supernatant was discarded and the gelatin microparticles were resuspended in a solution of 50 mM 4-(2-hydroxyethyl)-1-piperazineethanesulfonic acid (HEPES) (Corning) at pH 7.4. To remove the ethanol and Pluronic® F-127 the gelatin microparticle support bath was then washed three times with the same HEPES solution and stored until use at 4°C. Prior to printing, the uncompacted support was centrifuged at 1000 g for 3 minutes then washed with 50 mM HEPES. After the last wash, the gelatin microparticle support bath was suspended in 50 mM HEPES, degassed in a vacuum chamber for 15 minutes, and centrifuged at 1900-2100 g, depending on level of compaction desired, for 5 minutes. Finally, the supernatant was removed and the gelatin microparticle support bath was transferred into a print container.

### Collagen Bioink Preparation

All collagen type I bioink (LifeInk 200, Advanced Biomatrix). was prepared as previously described (*10*). Briefly, the stock 35 mg/mL LifeInk was mixed with syringes in a 2:1 ratio with 0.24 M acetic acid to produce a 23.33 mg/mL acidified collagen bioink. The bioink was then centrifuged at 3000 g for 5 minutes to remove bubbles. For printing, the collagen bioink was transferred to a 2.5 mL gastight syringe (Hamilton Company).

### Print Imaging and Visualization

Photographs of printed constructs were acquired using a Laowa 24 mm probe lens (Venus Optics) mounted to a Sony ILCE7M mirrorless digital camera. OCT 3D image stacks were acquired using a Thorlabs Vega 1300 nm OCT system with the OCT-LK4 objective (Thorlabs) (*27*). OCT images were prepared for visualization using Fiji (ImageJ, NIH) with noise reduction, median filtering, and stack histogram equalization. The image stacks were then exported as TIF files and opened using 3D Slicer (*30*). The volume rendering features were then used to produce 3D views of the volumetric OCT image.

### Print Gauging

Gauging was performed as we have previously described (*27*). Briefly, a 3D volumetric image of the ear construct was captured using OCT. The image was then cleaned and segmented to produce a 3D reconstruction using 3D Slicer (*30*). The 3D reconstruction and the original 3D model were then imported into CloudCompare (www.cloudcompare.com) (*31*). In CloudCompare the two 3D objects were aligned and registered and then the reconstruction was gauged to the original 3D model to determine errors. Errors were calculated as mean ± STD.

### Statistics and Data Analysis

Statistical and graphical analyses were performed using Prism 9 (GraphPad) and Excel (Microsoft). Statistical tests were chosen based on the experimental sample size, distribution, and data requirements. Analysis of the 1000 and 500 µm grid print was completed using Fiji (Image J NIH) and MATLAB (Mathworks). For the comparison of the two grid sizes a Student’s two-tailed unpaired t test was used.

## Supporting information

Supplemental Information - Conversion Guide

## Acknowledgments

We would like to acknowledge Brian Coffin and Andrew Hudson for reviewing the supplemental materials. We would also like to acknowledge the attendees of the Regenerative Biomaterials and Therapeutics Group’s 2021 3D bioprinting open-source workshop for real world testing of the conversion process. Additionally we would like to acknowledge Thingiverse (www.thingiverse.com) user ChrisGilleti, who designed the original model for the Duet2 WiFi Case utilized in this conversion (thing 3721923).

## Funding

This work was supported by the Food & Drug Administration (R01FD006582) and the National Heart, Lung, And Blood Institute of the National Institutes of Health (1F30HL154728, K99HL155777).

## Author contributions

All authors conceived the experiments and contributed to the scientific planning and discussions. J.W.T and D.J.S. prepared final figures and text. J.W.T wrote the conversion guide. J.W.T performed bioprinting and OCT imaging. J.W.T and D. J. S performed image analysis in Fiji and MATLAB. J.W.T, D.J.S., and A.W.F. wrote the paper and interpreted the data.

## Competing interests

A.W.F. has an equity stake in FluidForm Inc., which is a startup company commercializing FRESH 3D printing. FRESH 3D printing is the subject of patent protection including U.S. Patent 10,150,258 and provisional patent No. 63/082621.

## Data and materials availability

All digital models are available for download at (**ZENODO**). Any raw data not presented in the main and supplemental text is available on request.

## Notes

https://doi.org/10.5281/zenodo.7067499

https://doi.org/10.5281/zenodo.4119127

