## Supplemental Information - Conversion Guide for "Development of a High-Performance Open-Source 3D Bioprinter"

### Title

### 3D BIOPRINTER CONVERSION GUIDE

#### INTRODUCTION

This guide will show the reader how to convert a FlashForge Finder thermoplastic 3D printer into a bioprinter with a Duet3D Duet 2 WiFi control board and our Replistruder 4 syringe pump extruder. Where applicable we will describe the general process of converting a plastic printer to a bioprinter so that the reader can apply these principles to converting the printer of their choice (e.g., if the FlashForge Finder does not meet their needs).

#### REQUIRED COMPONENTS/TOOLS AND COST

| Component | Supplier | Part Number | Cost |
|---|---|---|---|
| FlashForge Finder (includes small toolset with necessary hex wrenches and Philips screwdriver) | www.amazon.com | B016R9E7J2 | $299.00 |
| Crimping Tool, DuPont Connectors | www.amazon.com | B08ZS22MZ9 | $33.99 |
| Fine Wire Stripper | www.amazon.com | B000XEUPMQ | $19.97 |
| Aviation Shears | www.amazon.com | B00OCGQFQQ | $14.99 |
| Precision Screwdriver Set | www.amazon.com | B0747DYJJR | $7.99 |
| Duet 2 WiFi | www.matterhackers.com | M3CWNU4L | $155.00 |
| Roll of PLA | www.matterhackers.com | MY6C8H7E | $42.00 |
| Replistruder 4 | https://doi.org/10.1016/j.ohx.2020.e00170 | N/A | $150.00 |
| From Replistruder 4 Build:<br>M3 Thin Nut<br>M3 x 20 mm Socket Cap Bolt | N/A | N/A | N/A |
| | | | **Total** |
| | | | $722.94 |

#### PRINTED COMPONENTS

| Component | URL |
|---|---|
| Duet 2 WiFi Case | **Add Zenodo Link** |
| FlashForge Finder X Axis Carriage | **Add Zenodo Link** |
| FlashForge Finder 35 mm dish holder | **Add Zenodo Link** |
| FlashForge Finder well plate holder | **Add Zenodo Link** |

### PRE-CONVERSION CONSIDERATIONS

#### Selection of a Plastic Printer to Convert

It is possible to convert a wide range of stepper-motor-driven 3D thermoplastic printers into a bioprinter. While there are many different configurations for cartesian plastic printers, some are better than others for bioprinting. In general, the ideal solution is a gantry where the X, Y, and Z axes can move the syringe pump without moving the sample at all. However, there are not many plastic 3D printers that use this configuration. The best alternative is a printer that only moves the sample in the Z axis, while the X and Y axes can move the syringe extruder without the sample being shifted. This is a common configuration for plastic 3D printers, including the FlashForge Finder we use in this paper, and one we have published with multiple times (*1–4*). The other common configuration is for the sample to sit on the Y axis stage while the X and Z axes move the syringe extruder. For FRESH 3D bioprinting moving the sample rapidly on the Y axis runs the risk of inducing shear in the gelatin microparticle support bath, which could affect the print. Nevertheless, we have adapted multiple printers with this configuration, including the Printrbot Simple Metal, LulzBot Mini and LulzBot Mini 2, all with excellent results (*5*, *6*).

The next consideration that must be made is to the motion control system. The most common drive system for plastic printers is where the X and Y axes are driven by one or more belts, while the Z axis is driven using a leadscrew. In general, as we have shown in this manuscript, the accuracy of belted X and Y axes systems driven by stepper motors is around 100 μm. Most Z-axis leadscrews take advantage of gravity to minimize backlash in the system, which leaves accuracy to be determined by the quality of the screw itself. Among low-cost plastic printers this configuration is quite common. Higher cost does not always mean higher quality, however, as there is often little difference between the accuracy and precision of printers below $3000. To better utilize a larger budget, higher quality motion systems using precision lead screws or even precision ball screws can be built using standalone stages (e.g., IKO, Parker-Hannifin, Misumi, Anaheim Automation).

Finally, it is necessary to decide if an unassembled kit or preassembled printer is to be selected. In general, a preassembled printer is likely to be better for most use cases. This is because for assembly at a factory the manufacturer can invest in alignment jigs and measurement tools that will be used for assembly, ensuring a baseline performance out of the box. If a lab were to invest in these tools (e.g., right angle squares, precision flat surfaces, torque wrenches, calipers, gauges), or if it already has them, then it is possible to assemble a printer with better performance than the manufacturer, if time and care are taken.

**Selection of a Motion Control System**

There are many motion control systems for 3D printers, but currently high-performance 32-bit systems with quality stepper controllers (e.g., from Trinamic) are the best option, and here we use those developed by Duet3D. The Duet 2 WiFi provides quality motion control with up to 256x microstepping for 5 axes and a user-friendly interface with easily modified system configuration files. The Duet 2 WiFi can be accessed through a local network and can also host its own network, depending on the configuration. In addition to the Duet 2 WiFi, we have previously utilized the Duet Maestro and the Duet 3 with a standalone single board computer (e.g., Raspberry Pi 4) to offload the processing associated with hosting the user interface. All of these motion control boards can be successfully used in adapting a plastic printer to a bioprinter, and have the key advantage of being more customizable than the motion control boards that typically come stock on the printers.

**Selection of an Extrusion System**

More detail regarding bioprinter extrusion systems can be found in the Replistruder 4 publication (*7*). Briefly, the most common extrusion system utilized by commercial 3D bioprinters is pneumatic. While pneumatics work at a basic level, they can have issues extruding small volumes and stopping extrusion quickly, which are key for printing small, high-resolution constructs. It is particularly difficult for pneumatic extruders to pull fluid back into the syringe barrel through small gauge needles, a technique for limiting erroneous extrusion of material termed retraction. These limitations can lead to errors in the printed object, such as over or under extrusion, which require sophisticated and expensive pneumatic systems to overcome. For these reasons, we exclusively use positive displacement syringe-pump based extruders such as the Replistruder 4. We have also released designs for the Replistruder 3 (which is almost wholly plastic printed), and syringe pumps for large-scale (>60 mL ink volume) bioprinting (*5*, *6*). The Replistruder 4 is compatible with a wide range of syringes up to 25 mL and can be adapted to new syringes. The dispensing tips are easily exchanged, and we have used tips with inner diameters down to 20 μm and up to ~1 mm. Using the Replistruder 4, we routinely print structures with positive (printed ink) and negative (regions without printed ink) features on the order of 200 μm (*1*, *2*, *7*).

**Design of the X-Axis Carriage**

When designing a carriage to mount the syringe extruder there are several considerations based on the specific printer being used. The FlashForge Finder has the preferred configuration where the X and Y axes move the syringe extruder independent of the Z axis moving the print container. In this case, the plastic extruder is attached to the X axis and must be completely replaced with the newly designed X carriage to hold the syringe extruder and interface with the drive mechanism (the belt that moves the X-axis). When replacing this type of carriage there are physical measurements from the printer that must be designed for. Chief among these are the diameter of the linear rails, the spacing between the linear rails, the outer diameter of the bearings, the length of the bearings, the width of the belt, the vertical spacing between the top and bottom of the belt, and the position of the belt relative to the linear rails. These dimensions can be measured using digital calipers.

It is also necessary to determine the space provided for the syringe pump within the context of these measurements. For the Replistruder 4, which is 42 mm wide the possible orientations are to have this width oriented along the X-axis or the Y-axis. In the case of the FlashForge Finder, there is sufficient space between the belt and the front linear rail to fit the Replistruder 4 with its 42 mm width oriented along the Y-axis. This configuration allows the Replistruder 4 to be between the two linear rails, which will provide the stiffest construction and minimize mechanical oscillations during motion. If the syringe pump can fit between the rails, as is the case in the FlashForge Finder, then the next step is to determine the height of the mounting points. The Replistruder 4 is significantly taller than typical plastic printheads, and has a motor mounted near its top. To decrease instability due to the printhead being essentially an inverted pendulum, it is best to lower the center of gravity of the syringe pump relative to the height of the linear rails. However, when lowering the syringe pump in this manner the Z height of the build volume for the printer is decreased, as the end of the syringe is moving closer to the print bed. Once an appropriate height is selected for the mounting point, all the necessary dimensions for the X-axis carriage are defined.

If a printer with another configuration of the axes is to be converted, the main steps for designing a carriage for the syringe extruder are similar. If the rails the plastic print head are mounted on are vertically aligned instead of horizontally aligned, it is possible to mount the syringe pump with its 42 mm width either aligned with the X-axis or the Y-axis, depending on which orientation gives the largest work volume, maximizes rigidity of the system, and allows easy access to swap syringes. Finally, if the printer is already designed to have swappable printheads, such as the LulzBot Mini 2, it can be advantageous to copy the mounting hardware that has already been developed and adapt it to the syringe pump for the fastest conversion.

### CONVERSION

#### Initial Assessment of Plastic Printer:

The first steps here are to determine the voltage and current capabilities of the power supply that the plastic printer is provided with and to estimate the power requirements of the printer. The FlashForge Finder comes with a 24V supply capable of providing 2.71 amps and the X, Y, and Z stepper motors are 0.8 amps/phase (Fig. S1). In this conversion, all we need to run are the board, the X, Y, and Z stepper motors, and the Replistruder 4 motor. We will go over the current settings for each of these elements later, but based on these requirements the included power supply should be sufficient. If the reader is interested in converting another printer, which would require more power, they could purchase an appropriate power supply, we have had good results using Meanwell 24V power supplies.

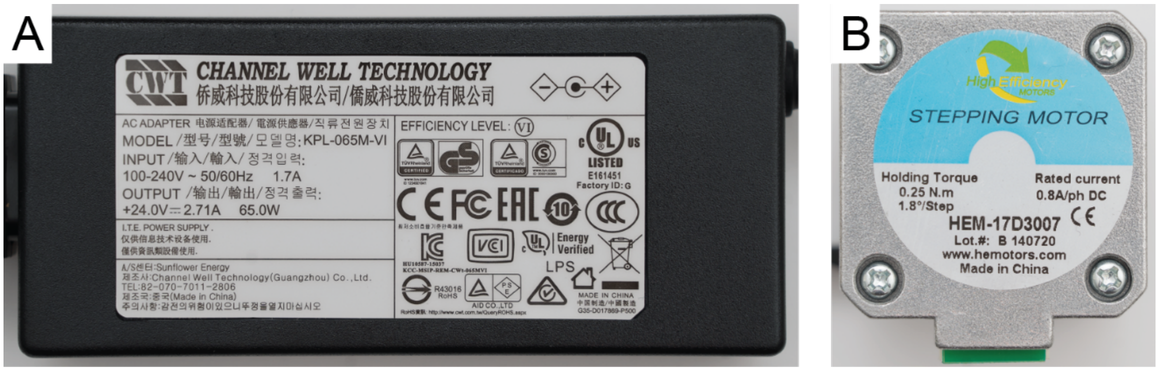


**Figure S1: Power Supply and Stepper Motors of FlashForge Finder**. (A) The included power supply can provide 2.71 amps at 24 volts. (B) The stepper motors driving the X, Y, and Z motion of the printer are 0.8 amps/phase.

The next step is to assess the electrical wiring and control board setup to begin to plan for the conversion. Prior to disassembling the printer make sure to shut it off and disconnect the power supply. The FlashForge Finder has its control board behind its back panel, which can be removed with four screws (Fig. S2A, blue circles). Make sure to keep track of these screws so you can put the back panel on again. Once the back panel is removed you can locate the control board (Fig. S2B, blue rectangle) and a fan (Fig. S2B, red rectangle). Taking a closer look at the control board, which in the case of the FlashForge Finder is well labelled, we can identify the relevant inputs and outputs for relocation to the new control board as well as some connections that we will not be using (Fig. S2C).

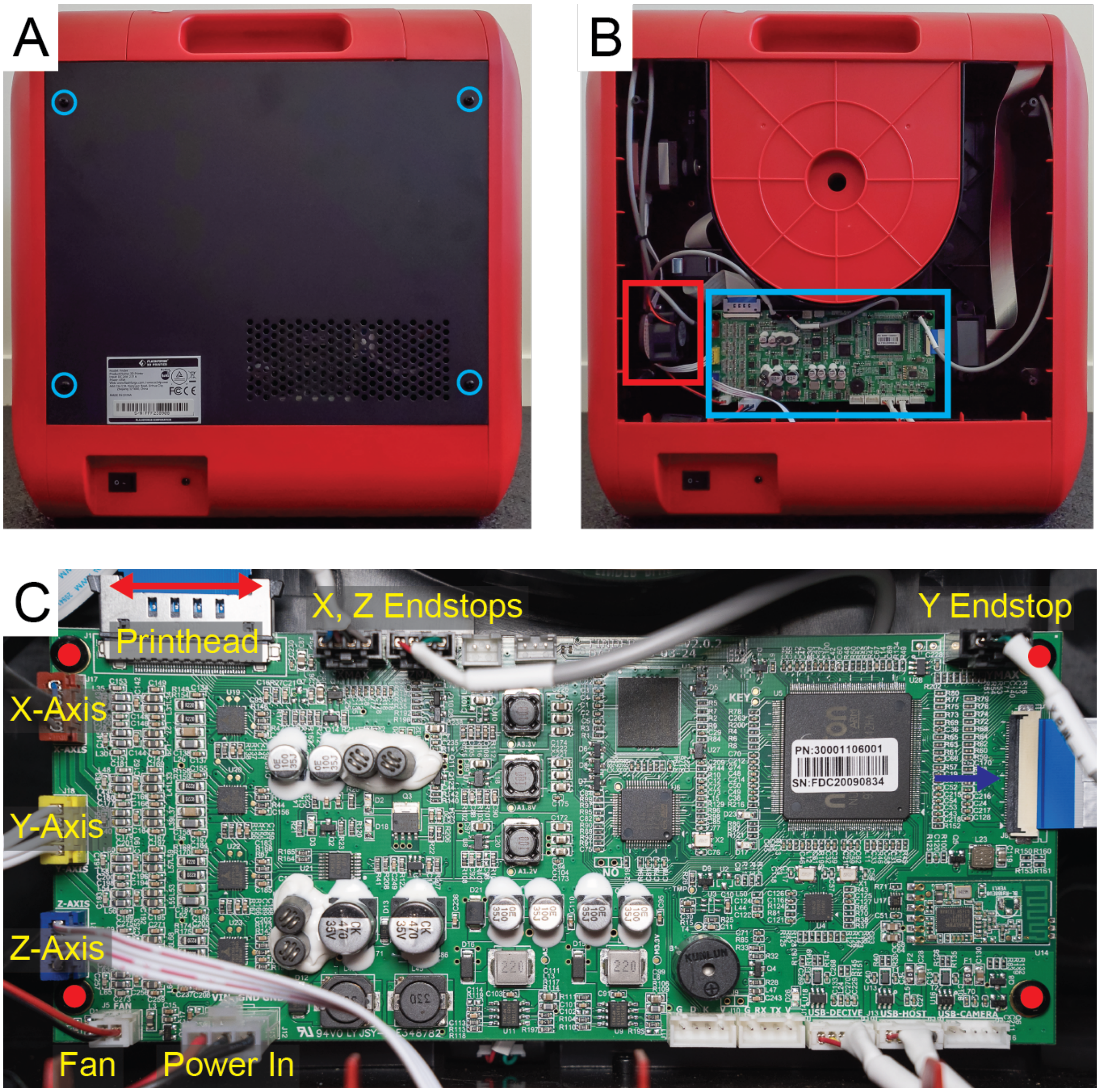


**Figure S2: Location of Control Board and Relevant Inputs and Outputs.** (A) For the FlashForge Finder the back panel can be removed using four black screws, highlighted in blue. (B) With the back panel off we can locate the control board (blue rectangle), the cables connected to it, and a 24V fan we will salvage (red rectangle). (C) Taking a closer look at the control board we can identify the relevant inputs and outputs as well as some that will not be utilized (red and blue arrows identify ribbon cable removal mechanisms, red dots identify mounting screws).

From this initial assessment we can tell that the power supply is sufficient for our needs and that the X, Y, and Z axes, their endstops, and the 24V fan are easily adapted to another board (they are easy to identify and well labelled). We won't be able to re-use the printhead connector as it is a proprietary ribbon cable, and we won't be able to utilize the display because it is not compatible with the Duet 2 WiFi.

The last connections on the board are USB port hookups and could likely be utilized on the Duet 2 WiFi, but this would require soldering and is beyond the scope of this conversion guide.

Next, we need to determine where to place the new control board. We want it to be somewhere easy to access and where the stepper, endstop, and power wiring can easily be rerouted. The FlashForge Finder has a built in plastic spool holder, which once removed leaves a space perfect for the Duet 2 WiFi (Fig. S3). To install the Duet 2 WiFi in this position we will need to find a case for it that fits, and we will need to extend the X, Y, and Z stepper motor cables to comfortably reach. For all non-extended wires, we will need to use the termination kit provided with the Duet 2 WiFi to change the terminations to mate with the board's connectors. We will also need to cut some small holes in the sides of the spool holder case to allow us to route these cables with the back panel reinstalled.

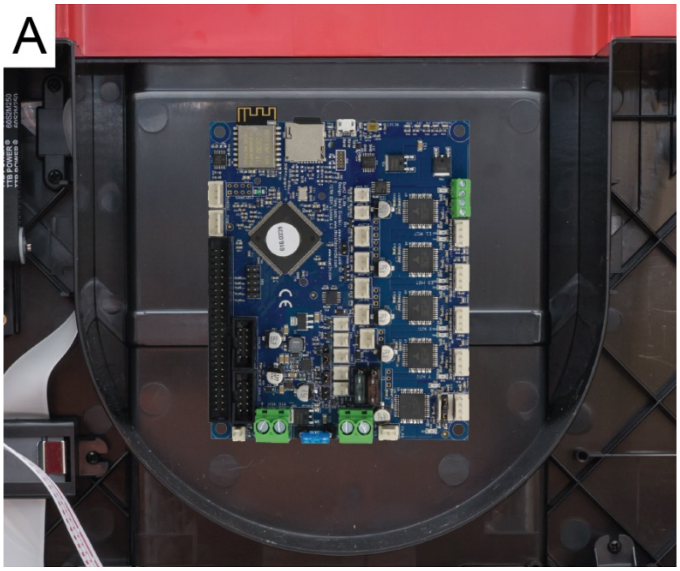


**Figure S3: Locating a Spot for the New Control Board.** A) The FlashForge Finder has a built in plastic spool holder, which when removed leaves a space that will fit the Duet 2 WiFi.

**Cleaning Up the Back Cabinet**

First, ensure the printer is not energized by disconnecting the power supply form the wall. Next, remove all the connections to the original control board. The printhead ribbon cable can be removed by squeezing the tabs on the side of the cable (Fig. S2C, red arrow). The display ribbon can be removed by flipping up the black clamp (Fig. S2C, blue arrow). After removing the board (by loosening 4 screws (Fig. S2C, red dots), you can salvage the 24V fan (Fig. S2B, red rectangle). To remove the 24V fan you can simply loosen the bolts attaching it to the printer (Fig. S4A, red circles). Keep these bolts, as we will use them to mount the fan elsewhere later. After removing the fan, we need to free the ribbon cable for the plastic printhead, so we can disassemble it later. To do this we need to remove two clamps (Fig. S4B).

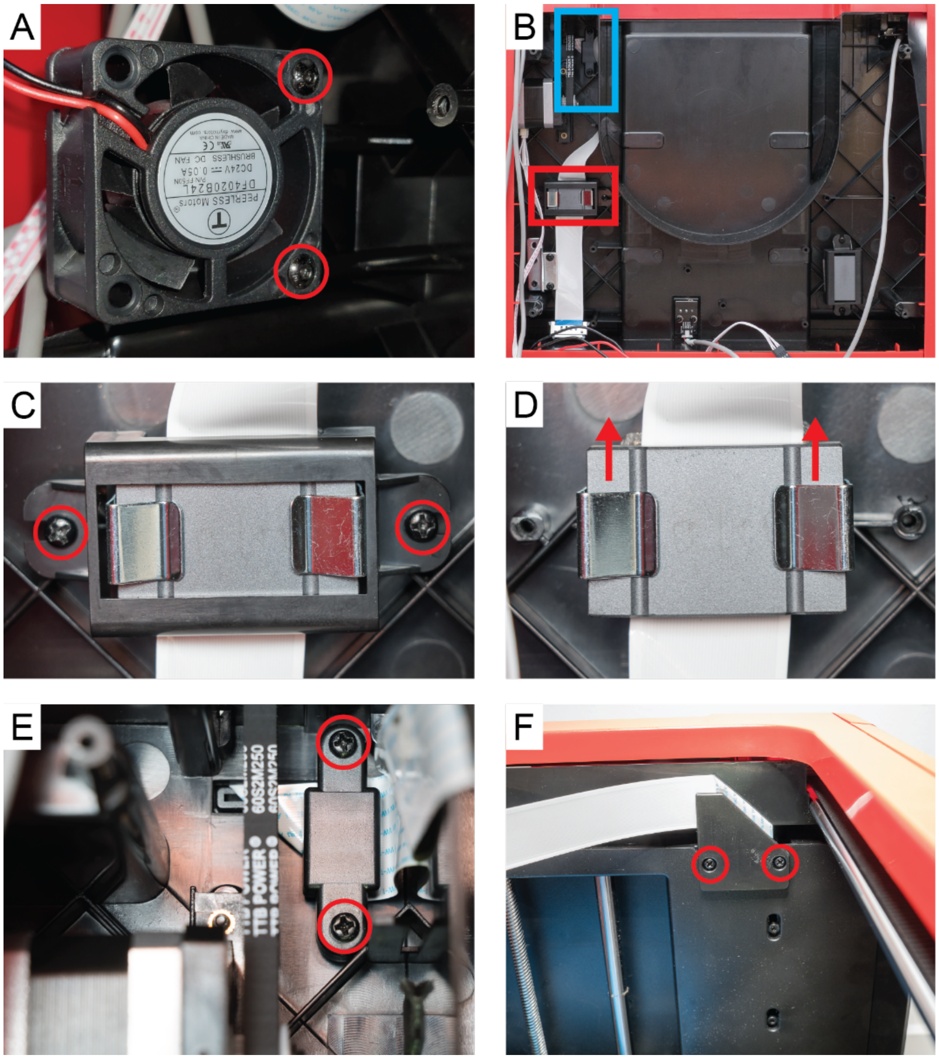


**Figure S4: Salvaging Fan and Removing Printhead Ribbon.** (A) The 24V fan can be removed by loosening the two bolts highlighted with red circles. (B) The ribbon cable is also anchored in the back cabinet with a ferrite (red rectangle) and a clamp (blue rectangle). (C) To remove the ferrite clamp, first loosen the two screws highlighted with the red circles and remove the plastic cover. (D) Next, pull the metal clamps apart and slide the top half of the ferrite upwards to free the ribbon cable. (E) The second clamp in the back cabinet can be removed by loosening the two screws highlighted with red circles. (F) At the front of the printer, to the right of the Z screw is the last clamp holding the ribbon cable. This can be removed by loosening the screws highlighted with red circles.

The first clamp is a ferrite clamp that can be removed by loosening two screws and sliding the ferrite away (Fig. S4, C and D). Keep these screws, as they will be useful in mounting the Duet 2 WiFi later. The second clamp is up near the Y axis stepper motor (Fig. S4B, blue rectangle). This clamp can also be removed by loosening the two highlighted screws (Fig. S4E, red circles). Finally at the front of the printer to the right of the Z screw the last clamp can be removed similarly to the previous one (Fig. S4F, red circles). After this the ribbon cable can be pulled through to the front of the printer.

**Preparation Prior to Installing the Duet 2 WiFi**

The first step to extending the X, Y, and Z stepper motor cables is to re-terminate them with a DuPont style connector. This process is demonstrated on the Z axis cable (Fig. S5 A to D). Prior to cutting the cables it is advisable to label them to be able to identify which axis they are for later. It is important to keep the cable wires in the same order, or to test for the two connected to each pole and keep the two pairs of poles in order when put in the 4 pin termination such that they are AABB. To test which wires connect to the same pole of the stepper motor you can take the cable to the step in Fig. S5D and touch two of the wires together then try to move the associated axis by hand. When the wires connected to the same pole are touching it will be harder to move the axis motor. The other two wires are by default the pair for the other pole of the stepper motor. This process can be utilized to ensure order in covered cables such as the X axis cable (Fig. S5F).

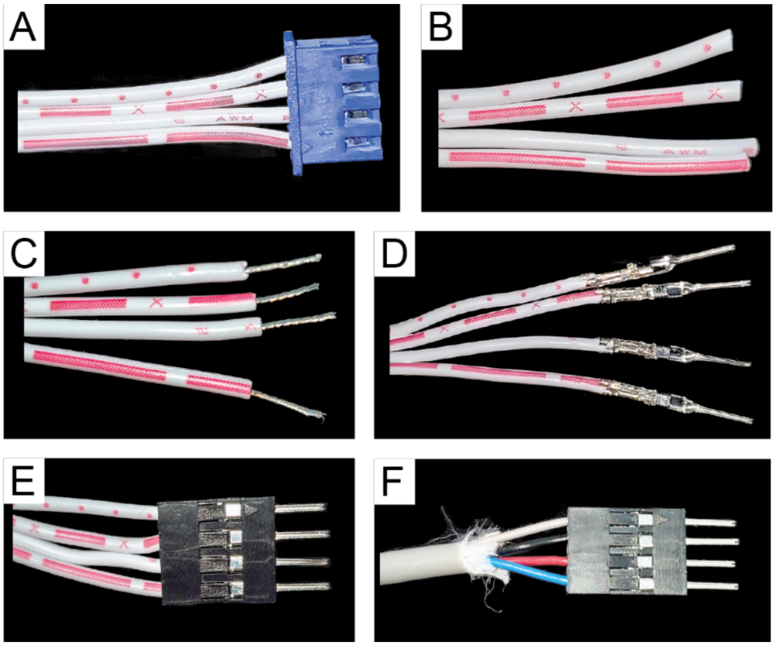


**Figure S5: Re-Terminating Stepper Motor Cables.** (A) The Z connector is initially not compatible with the Duet 2 WiFi. (B) The cable after trimming the termination off. (C) The cable after stripping 4-5 mm of the insulation off the individual wires. (D) The cable after crimping the individual wires with male DuPont 2.54 mm pitch connectors. (E) The Z connector cable with a 4 pin DuPont 2.54 mm pitch termination. (F) The X connector cable with a 4 pin DuPont 2.54 mm pitch termination.

Next, extensions for the re-terminated X, Y, and Z cables can be made. Here we have used 25 cm lengths of 4/22 shielded security system wire as it matches well with the wiring already found in the printer. On one side the process laid out previously (Fig. S5 A to E) should be repeated but utilizing female DuPont 2.54 mm pitch connectors (Fig. S6A). On the other side the process is very similar to Fig. 5 A to E, but the connectors provided by Duet 3D with the Duet 2 WiFi should be utilized, to be able to interface with the board (Fig. S6B). Here it is advisable to keep the individual wires in the same order to ensure that you can keep track of paired stepper motor poles.

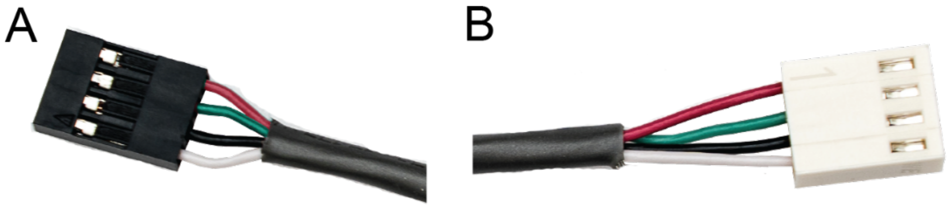


**Figure S6: Extenders for Stepper Motor Cables.** (A) A female 4 pin DuPont 2.54 mm pitch termination to accept the re-terminated stepper motor cables. (B) A female 4 pin connector provided by Duet3D with the Duet 2 WiFi.

The last three steps prior to installing the Duet 2 WiFi are to re-terminate the X, Y, and Z endstop cables, re-terminate the case fan (which we will salvage from the printer in this step) and to prepare the power cables. The endstops on the FlashForge Finder are connected to 4 wires, one of which is a redundant ground connection. The connections on the Duet 2 WiFi only accept a 3 pin connection, so we will need to identify the positive (VCC), ground (GND), and signal (SIG) wires.

The X, Y, and Z endstops are all slightly different, so it is necessary to inspect them to identify these three wires. The Y endstop can be found to the left of the Z stage (Fig. S7A). In our FlashForge the GND and VCC for this endstop were both, confusingly, black (Fig. S7B). To identify these wires on the other side of their cable we utilized a common multimeter, which can be set to identify two connected ends of a wire with a beep. The X endstop is located on the left side of the X axis (Fig. S7C). The Z endstop is visible after removing the back panel, in the lower middle of the back compartment (Fig. S7D). After determining the appropriate set of wires to adapt to the Duet WiFi connections each cable can be cut, stripped, crimped, and re-terminated with the Duet 2 Wifi connectors (Fig. S8 A to F).

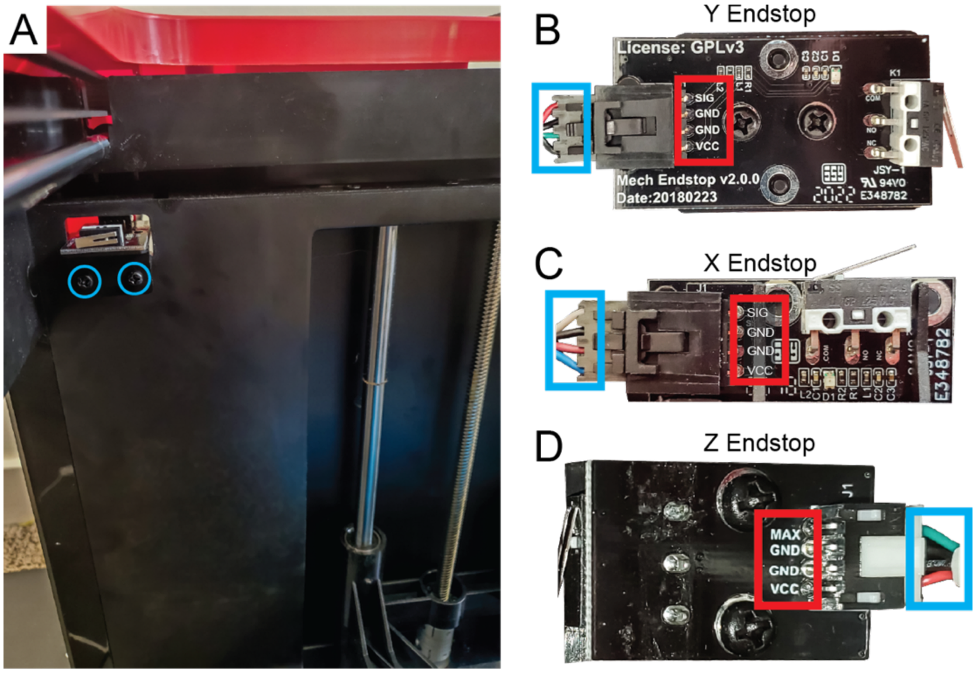


**Figure S7: FlashForge Finder Endstops.** (A) The Y endstop can be located left of the Z stage and can be inspected by removing the two screws highlighted by blue circles. (B) The four connections for the Y endstop are highlighted by the red box, their associated wires are highlighted by the blue box. (C) The four connections for the X endstop are highlighted by the red box, their associated wires are highlighted by the blue box. (D) The four connections for the Z endstop are highlighted by the red box, their associated wires are highlighted by the blue box.

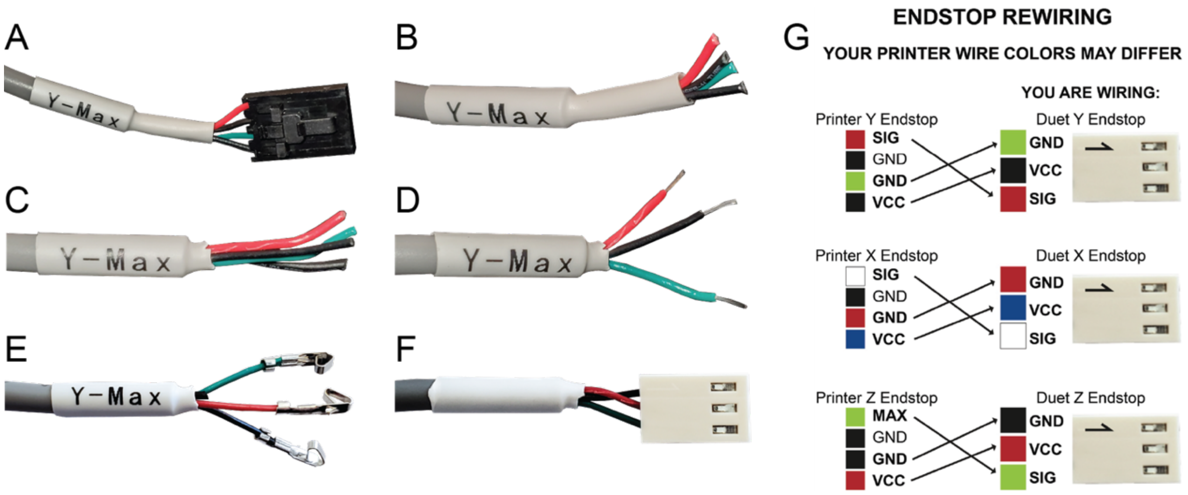


**Figure S8: Re-Terminating FlashForge Finder Endstops.** (A) The Y endstop cable terminated with the original connector. (B) The Y endstop with the original connector snipped off. (C) The Y endstop with all four wires exposed. (D) The GND, VCC, and SIG wires stripped for termination, with the fourth wire trimmed. (E) The GND, VCC, and SIG wires crimped with the Duet 2 WiFi female connectors. (F) The final terminated Y endstop, with the GND top, VCC middle, and SIG bottom, which matches the Duet 2 WiFi input. (G) An example endstop rewiring sequence for the Flashfordge Finder. Exact colors may differ between printers and should be checked with a multimeter.

After re-terminating the endstops we need to adapt the case fan originally used to cool the board that came with the FlashForge Finder (Fig. S9 A and B). To do this we simply need to trim the old connector off and re-terminate it with a 2 pin Duet 2 WiFi connector (as in Fig S8 A to F). We also need to prepare the power cabling. The FlashForge Finder's original power supply cable was terminated with 2 GND and one VCC wire. For the Duet 2 WiFi we only need one GND and one VCC. The Duet 2 WiFi has screw clamp connectors for VCC and GND in, so a bare wire can be inserted and a termination isn't required, though ferrules for such a termination are supplied by Duet3D. To adapt the FlashForge Finder cable we can simply trim the connector (Fig. S9 C and D). The remaining GND should be left in the connector and will later be contained with cable management. After trimming the power cable, the wires can be stripped and twisted (as in Fig S8D). The ferrite core that this cable is wrapped around must also be removed to provide sufficient cable length. With these preparations complete we have adapted all the original FlashForge Finder cables to the Duet 2 WiFi. Next, we can move to installing the control board itself.

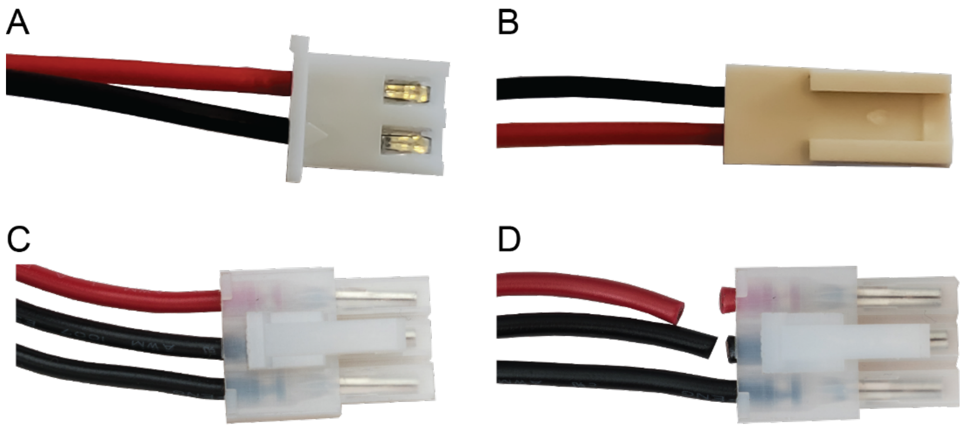


**Figure S9: Re-Terminating FlashForge Finder Fan and Power.** (A) The old fan connector. (B) The Duet 2 WiFi Fan connector. (C) The original power connection. (D) The power connection with VCC and GND separated.

**Configuring the Duet 2 WiFi**

For this step we recommend referring to the Duet 2 WiFi startup guide (guide) to get the most up to date instructions for your board. It is suggested that you update your firmware to the newest stable release; however, we recommend performing the firmware upgrade after the board set-up is complete and the Web Interface is running. To get the Web Interface running, you can decide to either connect your Duet 2 WiFi to an existing network (M587), or to set it up to host a standalone network to which you can connect directly (M589). We also recommend following the instructions for these configurations on the

Duet website. As an example, we will provide instructions that worked for our components to configure as a standalone network (*for Duet WiFi firmware older than 3.0 there might be some differences in the steps required for configuration). In our experience, for standalone network setup, we have found that the following series of commands through the terminal controller results in effective configuration in most instances. In some cases, both the Duet and Webserver firmware require updating before the standalone network will function properly depending on the age of your board and the firmware version preinstalled.

<u>Duet WiFi Start-up:</u>

1. Follow the guide at: https://duet3d.dozuki.com/Guide/1.)+Getting+Connected+to+your+Duet/7?lang=en
2. Launch the Windows (YAT) or Mac (SerialTools) program to connect to the printer board
    a. The board will appear as "usbmodem1411" or something similar at baud rate 115200
3. You will likely get a WiFi error due to no networks connected
    a. "WiFi reported error: no known networks found" this is normal and will be addressed

<u>Perform these commands after the WiFi startup guide has been completed:</u>

1. M552 S-1; Stops the WiFi Module
2. M115; Checks for the Duet2 WiFi Firmware. This will determine if you need to update later
3. M552 S0; Put WiFi card in idle mode
4. M588 S"*" ; This will clear any network SSIDs already present on the board
    a. *Mac users: Smart Quotes need to be disabled when sending G-code commands
        i. System Preferences>Keyboard>Text> Uncheck use smart quotes
5. M587 ; Checks to make sure no networks are listed
6. M589 S"PrinterNetworkName" P"Password" I192.168.0.1 C1 ;
    a. Adding the Network Name, Password, and IP address to the WiFi firmware.
    b. You can choose any network name and password, but some characters might cause issues so simple is better.
    c. *If error, the IP address might need to be changed. Increase incrementally (0.2 etc)
    d. For additional errors, refer to Duet web guide and help form.
7. M552 S2 ; Start WiFi card in access point mode.
    a. *This will need to be added to the config.g once connected to the control board.
    b. If another M552 command is present in the config.g, replace it with M552 S2.
8. Once the blue light on the WiFi module comes on, you are ready to connect to the hosted network
9. Launch a web browser and go to the IP address you defined in step #6

Once your Duet 2 WiFi is configured for access via the Duet Web Control, you can upload the files in the configuration folder we have provided (either the WiFi or Standalone versions) to the system directory (Fig. S10). Only upload the G-code file ".g". If you upload the system .bin files you can overwrite the firmware. If your firmware is older, you might need to upload the ".g" files to the sys folder on the SD card directly. Be careful if performing this step and refer to the Duet web guide and help documentation. The macro files we have provided can be uploaded under the macro tab at this point as well (Fig. S11). Finally, we suggest that you change the default jog distances for X, Y, Z, and the extruder (Fig. S12).

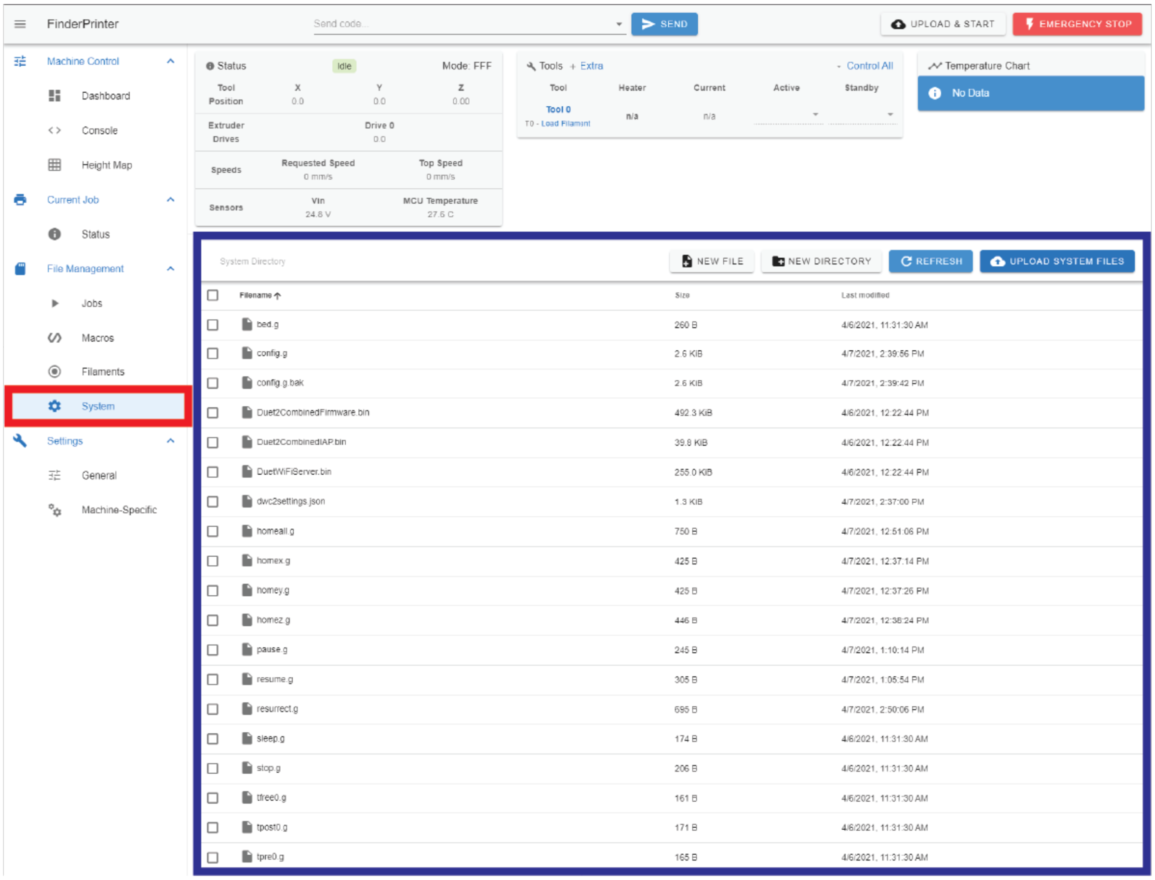


**Figure S10: Duet 2 WiFi System Editor.** The system editor can be found under the System tab (red rectangle) in Duet Web Control under File Management. Here you can edit the system files (blue rectangle).

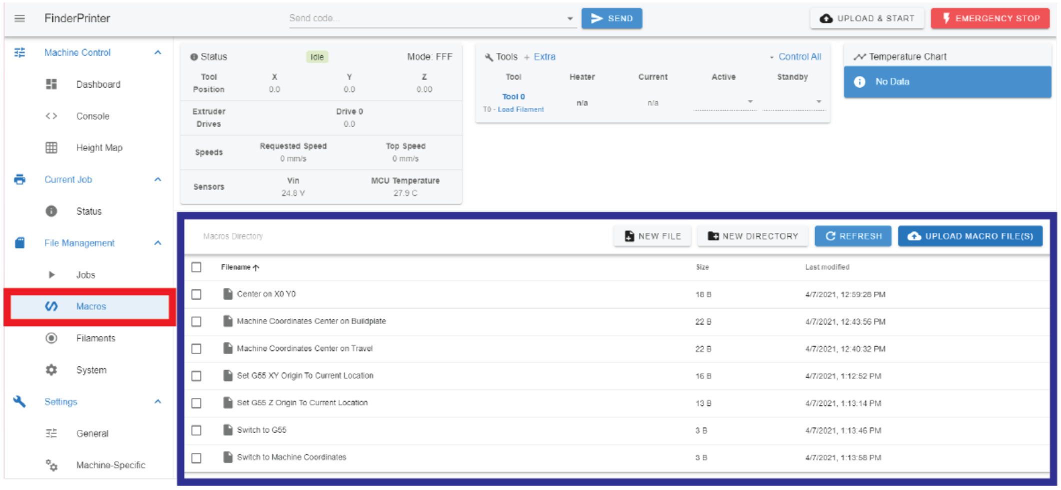


**Figure S11: Duet 2 WiFi Macro Editor.** The macro editor can be found under the Macros tab (red rectangle) in Duet Web Control under File Management. Here you can edit the macro files (blue rectangle).

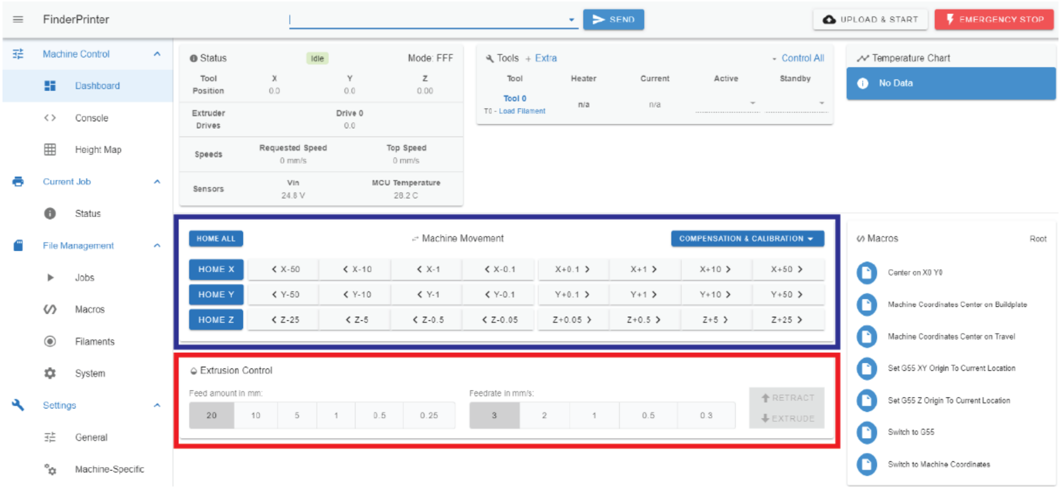


**Figure S12: Duet 2 WiFi Dashboard.** Here we suggest you change the jog distances under machine movement (blue rectangle) and the extrusion rate and distance under extrusion control (red rectangle).

#### Installing the Duet 2 WiFi onto the FlashForge Finder

Now that we have configured the Duet 2 WiFi we can install it onto the FlashForge Finder. First, we need to mount it to the case that we have already printed for it (Fig. S13A) using the screws that we salvaged when removing the printhead ribbon cables (Fig. S13B, red circles). We can also prepare to mount the case to the FlashForge Finder using double sided tape. First cut lengths of tape to fit the case (Fig. S13C, red lines). Next, adhere the tape to the case, making sure to leave the outward facing protective film intact (Fig. S13D). Prior to installing the case into the FlashForge Finder we need to slightly modify the walls of what used to be the filament spool holder. Use shears to cut the sides of 2 cm wide by 1.5 cm deep rectangles of plastic (Fig. S14A). Score the remaining attachment point of the plastic tabs and bend away from the score using pliers to snap the tabs out. After you have removed these tabs, install the Duet 2 WiFi and case onto the FlashForge Finder using the double sided tape previously applied (Fig. S14B).

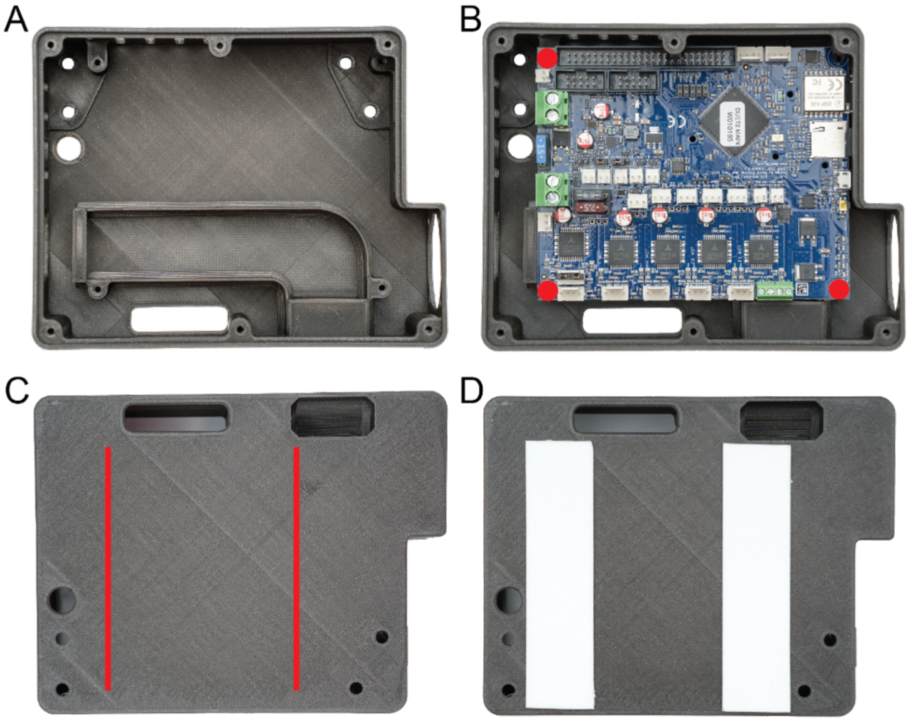


**Figure S13: Installing Duet 2 WiFi Into its Case.** (A) The case we have selected for the Duet 2 WiFi. (B) Mount the Duet 2 WiFi using the salvaged screws from the printhead ribbon cable clamps (3 screws, red circles). (C) Cut double sided tape to fit into the locations of the red lines. (D) Mount the double sided tape, leaving the outward facing protective film on.

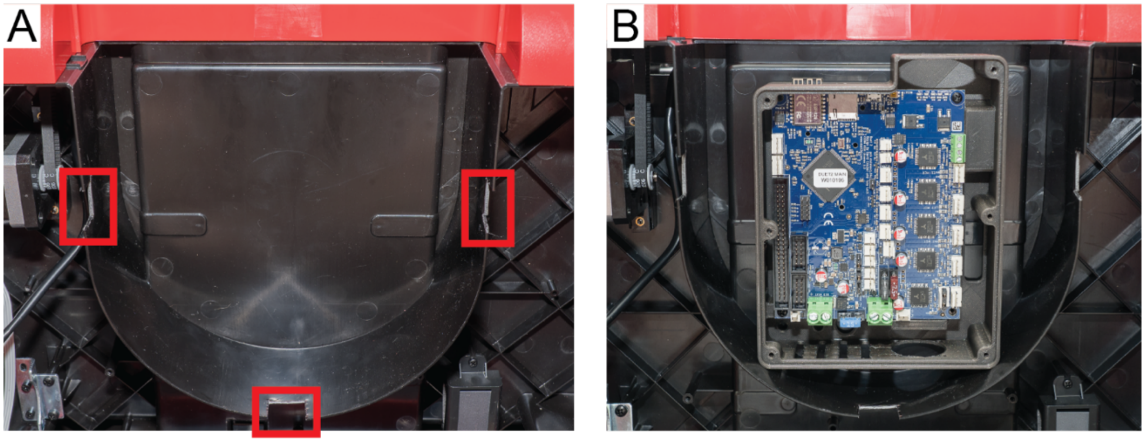


**Figure S14: Installing the Case onto the FlashForge Finder.** (A) Three tabs need to be cut away from the FlashForge Finder to allow for cable routing with the back panel installed (red rectangles). (B) The Duet 2 WiFi in its case can be adhered using the double sided tape in the orientation shown.

Finally, after attaching the Duet 2 WiFi and its case onto the FlashForge Finder we can complete the connections necessary to interface the board with the printer. First, connect the 3 extenders we have made for the X, Y, and Z stepper motors. Next, you can install the salvaged 24V fan at the top of the Duet 2 WiFi Case (Fig. S15A, blue rectangle). After this is complete, we defer to the Duet3D wiring guide (https://duet3d.dozuki.com/Guide/2.)+Wiring+your+Duet+2+WiFi-Ethernet/9?lang=en) noting that the 24V fan should be connected to one of the "always on" fan headers, and that we do not use any of the heating elements the board has to offer. After you have successfully connected the pre-prepared wiring, it is important to route the cables carefully (into the closest of the tabs we have cut) so that they remain attached with minimal strain on the connectors (Fig. S15, red rectangles, zip ties). The remaining wire and connector from the original power input can be zip tied into the lower bottom corner of the back cabinet, away from any metal. At this point we have successfully modified the cartesian motion components of the FlashForge Finder to be compatible with the Duet 2 WiFi and have successfully installed the board onto the FlashForge. We have also configured the Duet 2 WiFi to run these electrical components and host the Duet Web Control in the manner which is most appropriate (on an existing network or hosting its own network). The reader can now open Duet Web Control in a browser and home and then move the X, Y, and Z axes to verify that everything is correctly configured. If there is an issue you can refer to the Duet3D guides (https://duet3d.dozuki.com/c/Getting_Started) and forums (https://forum.duet3d.com).

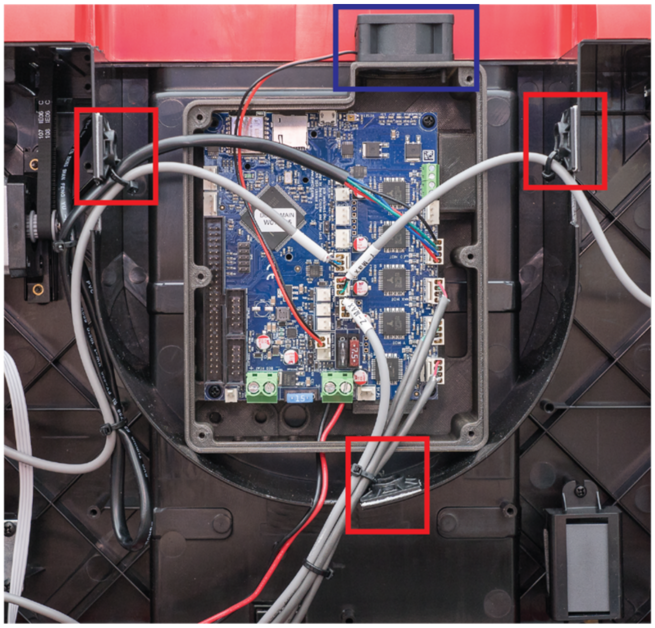

**Figure S15: Connecting the Duet 2 WiFi to the FlashForge Finder.** The salvaged 24V fan can be connected to the Duet 2 WiFi case (blue rectangle) using the bolts it was originally mounted with. Good cable routing is helpful, including zip-ties and zip-tie mounting bases (red rectangles).

**Modifying the X-axis to Hold the Replistruder 4**

Now that we have modified the FlashForge Finder to be controlled by the Duet 2 WiFi we need to remove the plastic printhead and replace it with a carriage that can hold a syringe pump, specifically the Replistruder 4, which we have detailed the construction of elsewhere. We have already begun this process by removing the ribbon connector earlier (Fig. S4 B to F). Next, we need to disassemble and remove the plastic printhead. To start we will disassemble the underside of the printhead, first removing the fan shroud and LED with three fasteners (Fig. S16 A and B). Next remove two more screws to aid in disassembly of the top of the printhead (Fig. S16C).

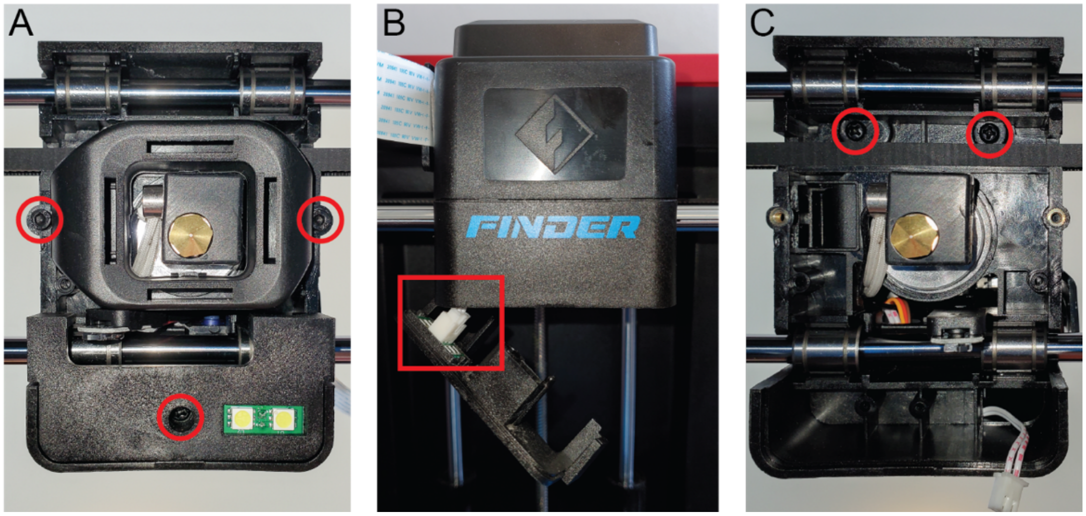


**Figure S16: Disassembling the Underside of the Plastic Printhead.** (A) Remove the screws and bolts highlighted with red circles. (B) Disconnect the LED panel. (C) Remove two additional screws, highlighted with red circles.

After disassembling the underside of the printhead, move to the top of the printhead to continue disassembly. Remove three bolts to reveal the printhead inner workings (Fig. S17A, red circles). Next, remove the ribbon cable (Fig. S17B, red arrow) and a single screw (Fig. S17B, red circle). Move to the left side of the printhead and remove an additional screw (Fig. S17C, red circle). Now move to the right side of the printhead and remove three more screws (Fig. S17D, red circles). This disassembly will allow you to remove the servo subassembly (translucent blue plastic piece) to reveal the green circuit board and more fasteners. Remove three more fasteners from the front of the green circuit board (Fig. S18A). This will release the hotend below the black plastic carriage and the circuitboard above (Fig. S18B). Cut the white heater cables, and black thermistor cables connecting these two pieces (Fig. S18B, red rectangles). Now all that remains is the black plastic carriage and the belt. The bearings on this carriage are well secured.

To removed them the general idea is to widen the bearing pockets by flexing the carriage (Fig. S19A) and then lifting the carriage off of the bearings. Before removing the bearings carefully cut the belt at one of the troughs of the teeth (Fig. S19B, red rectangle). Hold both ends of the belt as you do this to prevent it from slipping over the pully on the left side of the X axis (be careful not to lose this pully). Move the belt ends out of the way. The front bearings can be removed by hand, carefully pulling towards you (Fig. S19B, red arrows), and pushing off of the linear rods (Fig. S19B, yellow circles). For the rear bearings, a large flathead screwdriver must be used to disperse the force over the linear rod (Fig. S19C, red rectangles) and spread the bearing pockets (Fig. S19C, blue arrows).

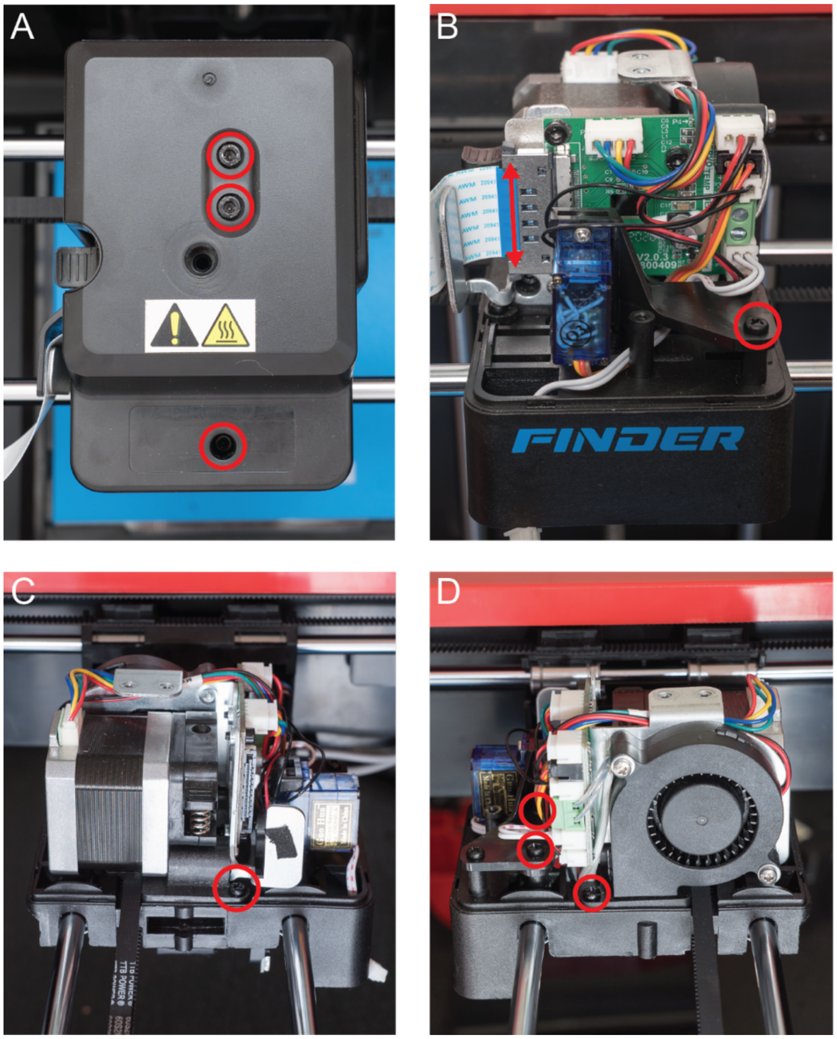


**Figure S17: Disassembling the Top of the Plastic Printhead.** (A) Remove the screws and bolts highlighted with red circles to reveal the printhead. (B) Disconnect the ribbon connector by squeezing the grey tabs (red arrow). Remove the screw highlighted by the red circle. (C) Remove an additional screw highlighted by a red circle from the left side of the printhead. (D) Remove three more screws, highlighted by red circles, from the right side of the printhead.

Carefully wiggle the bearings out of their pockets while applying force with the screwdriver. After you have successfully removed the front and rear bearings from the black plastic carriage we have completed disassembly and we can now begin to install the new carriage and the Replistruder 4.

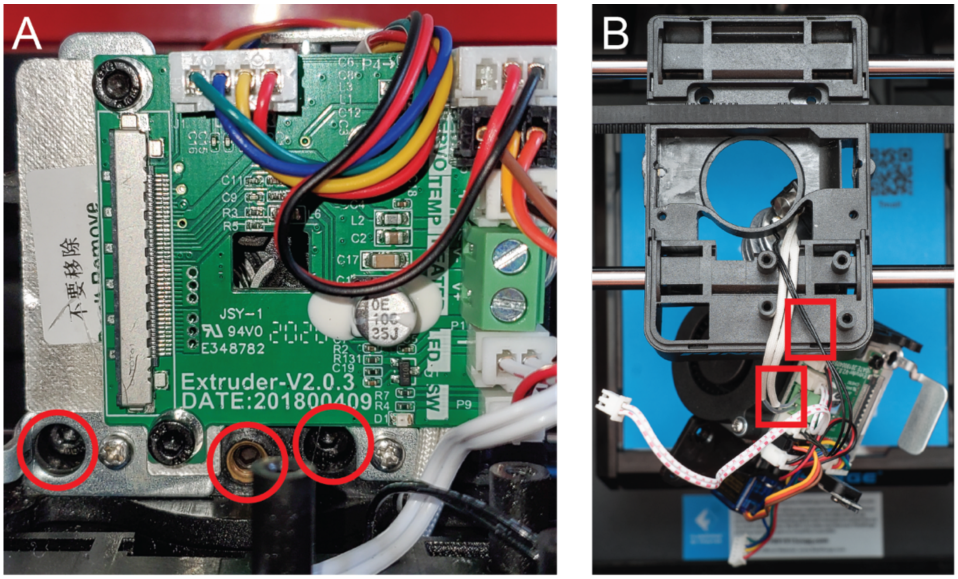


**Figure S18: Removing the Plastic Printhead.** (A) Remove three fasteners marked by the red circles. (B) The hotend and top assembly of the printhead will separate and be trapped by some cables, cut the hotend heater and thermistor cables (black and white cables attached to hotend) at the red rectangles to release.

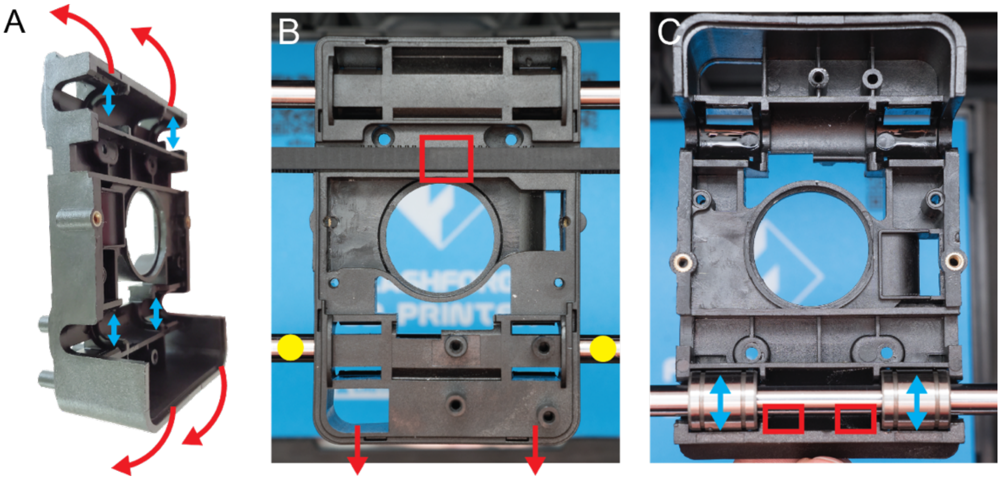


**Figure S19: Removing the Plastic Printhead Continued.** (A) The goal for removing the black carriage is to expand the bearing pockets (blue arrows) by creating a torque around the axis (red arrows). (B) Cut the belt in the trough of a tooth at red rectangle, holding both sides. Separate front bearings from the black plastic piece by carefully spreading the bearing pockets and pushing the black carriage up using your thumbs on the yellow circles. (C) Flip black plastic piece and remove rear bearings by carefully widening their pockets using a large flat head screwdriver.

**Installing the New Carriage and Replistruder 4**

Now that the entire plastic printhead has been removed we can install the new X axis carriage and the Replistruder 4 to complete assembly of our bioprinter. First, we will go over the installation of the carriage that we have designed, then we will discuss how to design a carriage for other printers. First, we need to assemble the new carriage. To trap the belt, we will use four M3 x 20 mm socket cap bolts (leftover from Replistruder 4 build) installed on the top of the carriage (Fig. S20A, red circles). Drive these bolts until the ends are just visible at the top of the upper belt channel (Fig. S20B, red rectangle) on either side of the carriage. Finally, install four M3 hex nuts (leftover from Replistruder 4 build) into the hexagonal pockets on the back mounting pillar of the carriage (Fig. S20C, red circles). These will be used to secure the Replistruder 4 to the carriage. You can pull the M3 nuts from the other side using an M3 x 20 mm bolt.

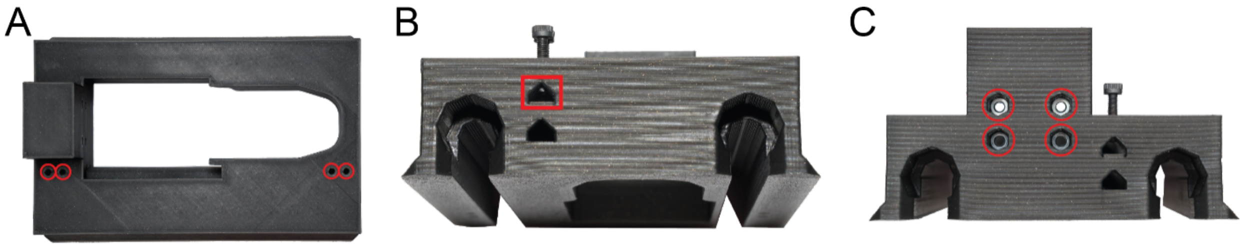


**Figure S20: Assembling the New Carriage. The (**A) Install four M3 x 20 mm socket cap bolts into the holes highlighted by the red circles. (B) Drive the M3 x 20 mm socket cap bolts until the ends are just showing on either side of the top belt channel (red rectangle). (C) Install four M3 nuts into the hexagonal pockets highlighted by the four red circles.

Next, assemble the Replistruder 4 according to our paper in HardwareX. The only slight deviation here will be that the stepper motor cable attachment point should be pointing to the left side of the Replistruder if you are looking at the front of the Replistruder. This will allow us later to route the cable so that it won’t bunch up as the carriage moves. Once the Replistruder 4 is assembled you can install it into the carriage. First, angle the Replistruder forward and slide the bottom into the open pocket in the carriage (Fig. S21A). Once the bottom is through, lower the Replistruder further into the pocket in the carriage (Fig. S21B). Finally, push the Replistruder back against the back mounting pillar and use four M3 x 20 mm socket cap bolts to mount the Replistruder to the carriage (Fig. S21C).

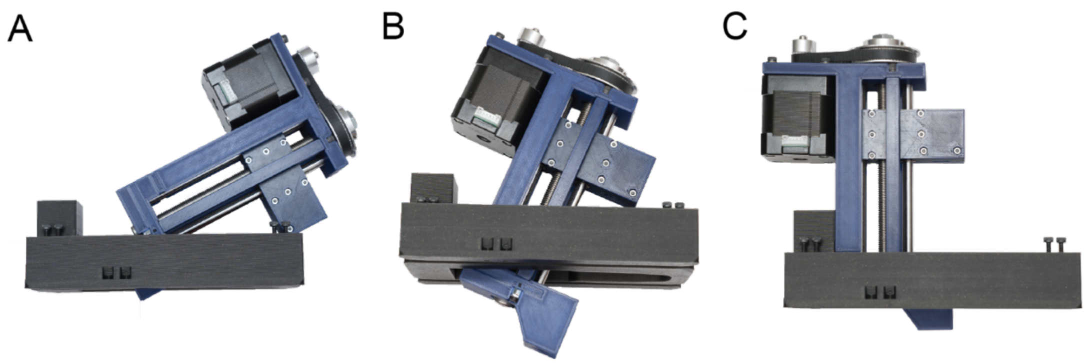


**Figure S21 Installing the Replistruder 4.** (A) Angle the Replistruder 4 forwards to fit into the pocket in the carriage. (B) Move the body of the Replistruder 4 lower into the carriage. (C) Push the Replistruder 4 backwards against the back mounting pillar (red rectangle), secure with four M3 x 20 mm socket cap bolts (through the bolts in Fig. 20C, red circles).

After assembling the carriage, we need to install it on the rails. To do so, spread the X axis bearings (Fig. S22A) and then place the carriage down such that the X axis linear rails fit into the two channels in the carriage (Fig. S22B). Next, on the left side, insert the bearings into their pockets (Fig. S22C, blue arrows). Thread the belt, tooth side down, into the left top belt channel (Fig. S22C, red rectangle) until it is visible next to the Replistruder (Fig. S22C, yellow arrow) and tighten the M3 x 20 mm socket cap bolts until just tight (Fig. S22C, red rectangle). On the left side of the X axis loop the belt around the pully (Fig. S22 D, red rectangle) and thread it into the left side lower belt channel (Fig. S22E, red arrow). Bring the belt all the way to the right side of the X axis and loop it around the motor pully (Fig. S22F, red rectangle). Insert the right bearings into their pockets (Fig. S22G, blue arrows). Next, thread the belt into the right top belt channel (Fig. S22G, red arrow). Finally, pull the belt until taut with needle nose pliers (Fig, S22H, red rectangle), then tighten the M3 x 20 mm socket cap bolts until just tight (Fig. S22H, blue rectangle). The carriage is now mounted to the X axis rails and the belt. Make sure that the bearings on both sides are pushed all the way into their pockets.

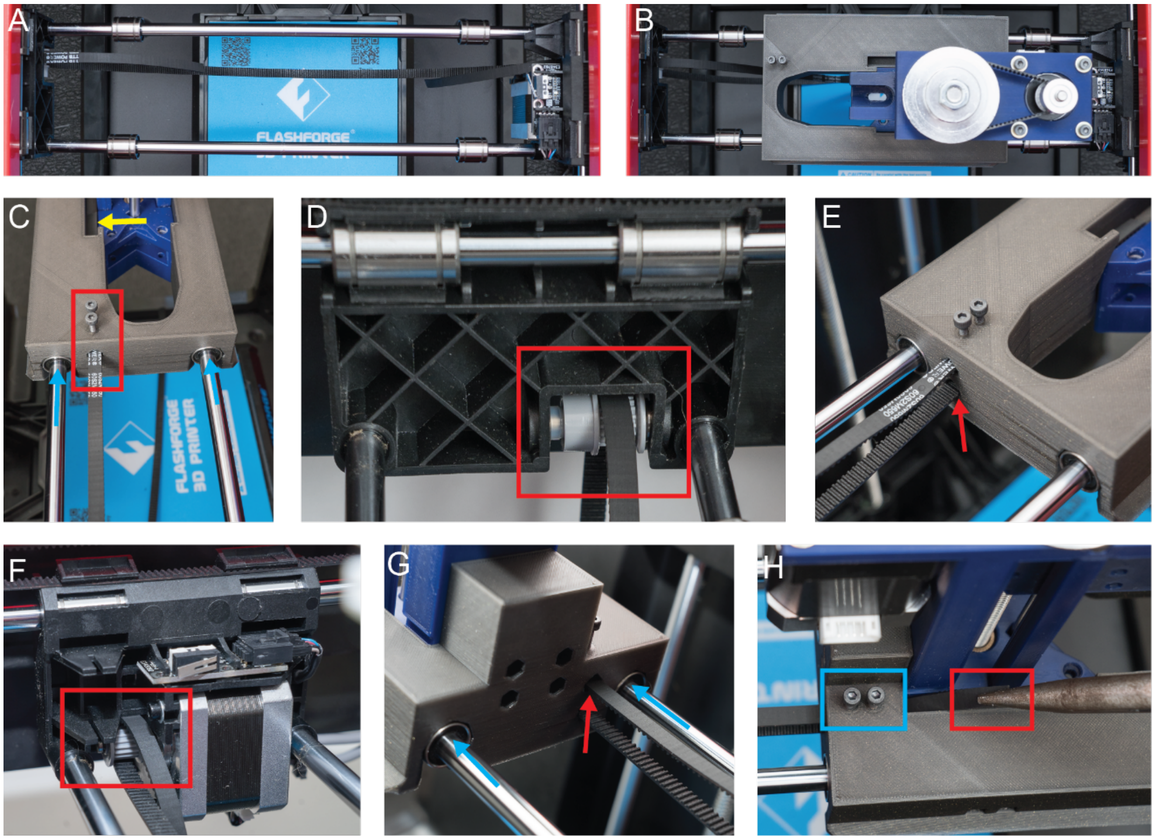


**Figure S22: Mounting the Carriage on the Rails.** (A) Prior to placing the carriage on the rails push the bearings apart. (B) Place the carriage between the bearings, ensuring the rails go into the two channels in the carriage. (C) Push the left side bearings into their pockets (blue arrows). Thread the top belt, tooth side down, into the left side of the top belt channel until the tip can be seen next to the Replistruder (yellow arrow). Tighten the M3 x 20 mm socket cap bolts until they grip the belt well (red rectangle). (D) Loop the belt tooth side down around the left side pully (red rectangle, not attached to motor). (E) Thread the bottom belt, tooth side up, into the left side of the bottom belt channel (red arrow), all the way to the other side of the X gantry. (F) Loop the belt, tooth side up around the motor pulley (red rectangle, attached to motor). (G) Insert the right side bearings into their pockets (blue arrows). Thread the top belt, tooth side down, into the right top belt channel (red arrow). (H) Pull the top belt using needle nose pliers (red rectangle) just until taut and tighten the M3 x 20 mm socket cap bolts until they grip the belt well (blue rectangle).

#### Finalizing Connections and Installing a Syringe

After mounting the Replistruder 4 the stepper motor cable must be connected to the Duet 2 WiFi. The cable can be passed through the port where the old plastic printhead's ribbon cable emerged (Fig. S23A, blue rectangle). At this point you can power down the Duet 2 WiFi and connect the Replistruder's stepper motor cable to the fourth stepper motor driver, E0. It is advisable to also incorporate the Replistruder's cable into your cable management in the back cabinet of the FlashForge Finder. The back cabinet can now be closed, taking care to ensure the cables are not pinched, using the original screws saved during disassembly.

A syringe, in its appropriate clamp adapter, can now be loaded into the Replistruder 4 (Fig. S23B). The syringe is first passed into the space in front of the Replistruder 4 (Fig. S23C). It is then inserted into the Replistruder 4 and secured using the appropriate bolts (Fig. S23 D and E). To ensure that the stepper motor cable is not disturbed during printing it is also good practice to secure a 25 cm length of cable using a zip-tie mount (Fig. S23F). At this point the FlashForge Finder conversion to a bioprinter with a Replistruder 4 is complete.

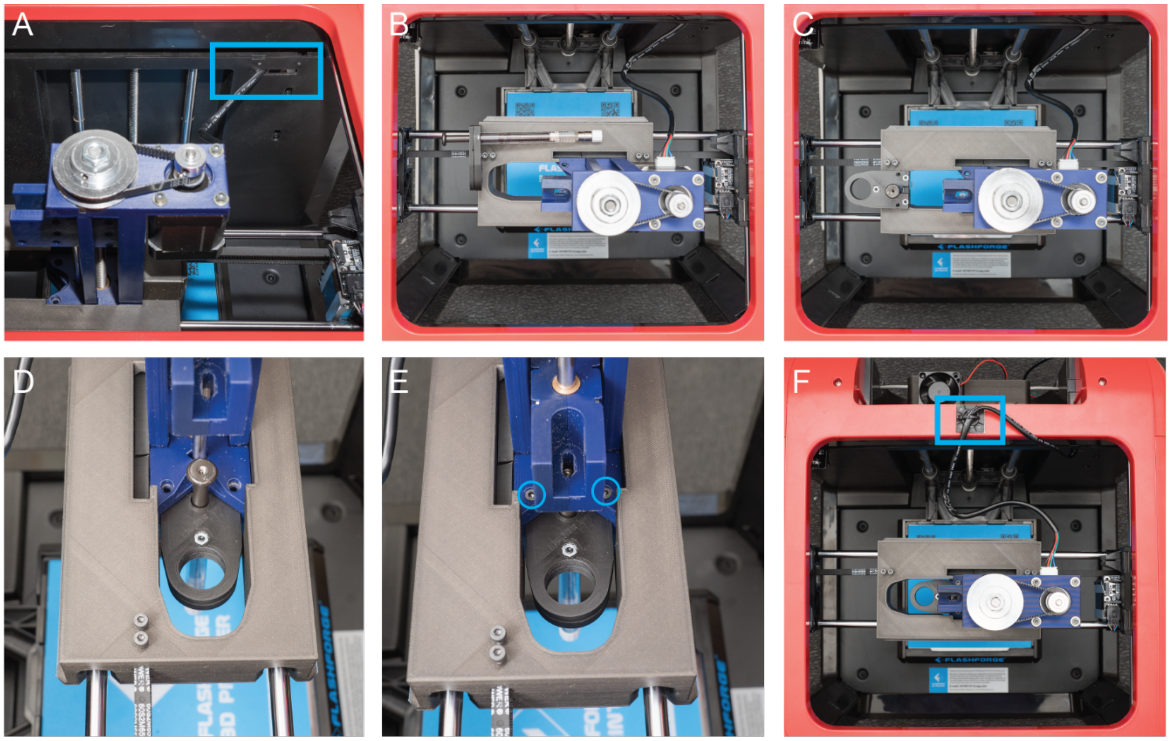


**Figure S23: Wiring the Replistruder 4 and Mounting a Syringe.** (A) After connecting the Replistruder 4's stepper motor cable it can be threaded through the port where the old ribbon cable used to emerge (blue rectangle). (B) When loading a syringe, it is first clamped into its appropriate size adapter. (C) The syringe is then passed through the space in front of the Replistruder 4. (D) The syringe adapter can then be inserted into the Replistruder 4. (E) The adapter can then be secured to the Replistruder 4 with the appropriate bolts. (F) After completing wiring, it is again advisable to place a zip-tie mount (blue rectangle) and secure a 25 cm free length of stepper motor cable.

**Build Platform Trays**

Two additional components that can be 3D plastic printed have been designed to aid in the bioprinting process. The first of these is a tray for the build platform that will hold a 35 mm petri dish, which is a common printing vessel. The second is a tray that will hold a well plate, which can be used for printing multiple constructs. The files for these trays are available for download.

**Process for Initializing a Print Using the Bioprinter**

With the bioprinter conversion process complete, the next step is to 3D bioprint a construct. While this bioprinter can be used with a wide array of bioprinting methodologies, we will describe the process for FRESH 3D Bioprinting using collagen type I. The first step is to prepare G-code for controlling the printer. To do this we utilize the open-source 3D slicing program Slic3r (slic3r.org), for which we have provided a configuration (.ini) file. This configuration file is designed for collagen type I in a 2.5 mL Hamilton Gastight syringe terminated with a 150 µm ID needle tip. After importing the configuration into Slic3r, load the construct to print and center it at the origin of the build plate. Using the provided settings, export the G-code. Next, prepare the FRESH gelatin microparticle support bath and collagen bioink as described in the main text methods section. To hold the gelatin microparticle support bath we will utilize a 35 mm petri dish and the provided 35 mm petri dish tray.

Turn the 3D bioprinter on and establish a connection to the Duet Web Control. Upload the G-code file prepared earlier. With the collagen bioink loaded in the syringe pump and the dish filled with gelatin microparticle support bath (held in the custom tray) we need to define the coordinate system of the dish. To do this, first home all of the printer axes using the "Home All" button on the interface. Next, use the X and Y axes of the "Machine Movement" section of the interface to jog the needle of the syringe to roughly the center of the dish. Now, use the Z axis of the "Machine Movement" section of the interface to carefully jog the needle of the syringe to be just slightly above the bottom of the 35 mm petri dish filled with gelatin microparticle support bath. At this point, the tip of the needle is at the center of the bottom of the print. Click the "Set G55 XY Origin To Current Location" macro on the Duet Web Control. Also click the "Set G55 Z Origin To Current Location" and "Switch to G55" macros. At this point we have defined the center of the G55 coordinate system as the current location of the needle tip and have switched the printer to the G55 coordinate system. Initiate the print process by selecting the previously uploaded G-code in the "Jobs" tab under "File Management" on the Duet Web Control. Once printing is complete, the gelatin microparticle support bath can be melted and washed away, after which the printed construct can be retrieved.
